# Simultaneous Profiling of Chromatin Accessibility and DNA Methylation in Complete Plant Genomes Using Long-Read Sequencing

**DOI:** 10.1101/2023.11.15.567180

**Authors:** Basile Leduque, Alejandro Edera, Clementine Vitte, Leandro Quadrana

## Abstract

Epigenetic regulations, including chromatin accessibility, nucleosome positioning, and DNA methylation intricately shape genome function. However, current chromatin profiling techniques relying on short-read sequencing technologies fail to characterise highly repetitive genomic regions and cannot detect multiple chromatin features simultaneously. Here, we performed Simultaneous Accessibility and DNA Methylation Sequencing (SAM-seq) of purified plant nuclei. Thanks to the use of long-read nanopore sequencing, SAM-seq enables high-resolution profiling of m6A-tagged chromatin accessibility together with endogenous cytosine methylation in plants. Analysis of naked genomic DNA revealed significant sequence preference biases of m6A-MTases, controllable through a normalisation step. By applying SAM-seq to Arabidopsis and maize nuclei we obtained fine-grained accessibility and DNA methylation landscapes genome-wide. We uncovered crosstalk between chromatin accessibility and DNA methylation within nucleosomes of genes, TEs, and centromeric repeats. SAM-seq also detects DNA footprints over cis-regulatory regions. Furthermore, using the single-molecule information provided by SAM-seq we identified extensive cellular heterogeneity at chromatin domains with antagonistic chromatin marks, suggesting that bivalency reflects cell-specific regulations. SAM-seq is a powerful approach to simultaneously study multiple epigenetic features over unique and repetitive sequences, opening new opportunities for the investigation of epigenetic mechanisms.

## BACKGROUND

Chromatin is a highly dynamic structure that plays a pivotal role in the regulation of eukaryotic gene expression. It is primarily structured as arrays of nucleosomes, which are separated by DNA segments of variable size often containing regulatory sequences. DNA wrapped into nucleosomes is generally inaccessible to other proteins. Thus, changes in the density and localization of nucleosome arrays provide a layer of epigenetic regulation (1). DNA itself can also carry epigenetic modifications, notably in the form of cytosine methylation (5mC), which is a repressive mark associated with the silencing of transposable elements (TEs) and genes in both plants and mammals (2). How nucleosome organisation and DNA methylation affect each other and the regulation of gene activity is still not fully understood. This lack of knowledge is partly due to the inability of current technologies to simultaneously chart both DNA methylation and chromatin accessibility across strings of nucleosomes (3). Indeed, techniques such as MNase-seq dedicated to map nucleosome positioning, ATAC-seq to chromatin accessibility, and BS-seq to DNA methylation, rely on short-read sequencing technologies spanning fragments of less than one nucleosome in size. Furthermore, due to their short length (<150 nt-long), these sequencing reads cannot be aligned unambiguously over highly repetitive sequences, such as centromeres, telomeres, and high copy TEs, thus providing an incomplete characterization of chromatin regulation. Long-read sequencing technologies, such as Oxford Nanopore Technology (ONT), have overcome this limitation by reading long DNA molecules while simultaneously detecting base modifications. The ability of ONT to sequence native DNA also avoids amplification biases, leading to improved estimation of DNA modifications (4). Recent works have exploited these characteristics to study DNA methylation together with chromatin accessibility in different species (5–8). For instance, it is possible to treat chromatin with bacterial m6A methyltransferases (m6A-MTases), which preferentially methylate accessible adenines due to steric hindrance. Because of the lack of endogenous m6A in most eukaryotes, treatment of chromatin with m6A-MTases followed by ONT sequencing could provide a high-resolution snapshot of chromatin accessibility, as was demonstrated using the single-molecule long-read accessible chromatin mapping (SMAC-seq) sequencing assay in yeast and humans (9). More recently, targeted analysis of centromeres, telomeres, and rDNAs in the model plant species *Arabidopsis thaliana* has been reported (10), opening the possibility to implement m6A-MTases-based approaches to study chromatin in plants. However, broader use of this method for simultaneous analysis of chromatin accessibility and DNA methylation on a genome-wide scale in Arabidopsis and other plant species is yet to be explored. Moreover, its potential to unravel the intricacies of large and complex genomes, such as those of crops, remains to be validated. Crucially, DNA modifying enzymes are known to have sequence preferences, which must be taken into consideration when used to assess chromatin landscapes (11, 12), even more importantly when tracking chromatin modifications at the genome-wide scale. Whether M6A-MTases have sizable sequence preferences remains elusive, and further characterizations need to be performed to ascertain the validity of results obtained with these approaches (5, 9, 10).

Here, we adapted SMAC-seq (9) to study high-resolution chromatin profiles of plant genomes. The simultaneous accessibility and DNA methylation sequencing (SAM-seq) data obtained here enabled us to study multiple epigenetic features in Arabidopsis and maize nuclei. We found that m6A-MTases have significant DNA substrate preferences, leading to sequence- and strand-biased m6A levels. We show that normalisation of m6A levels is critical to obtain reliable results, especially for the analysis of highly repetitive sequences such as Arabidopsis centromeric repeats. We demonstrate that SAM-seq can be used to identify large scale chromatin compartments, discriminate active and repressed regions, obtain high-resolution profile of nucleosome positioning and DNA methylation, identify DNA footprints at occupied transcription factors binding sites, and map chromatin landscapes across lowly and highly repetitive sequences.

## MATERIALS AND METHODS

### Plant material

Arabidopsis thaliana seeds (Col-0 accession) were germinated and grown in long-days (16h:8h light:dark) conditions at 23°C for approximately three weeks. Rosette leaves were harvested, flash frozen in liquid nitrogen, and stored at −80°C until use. Maize seeds (B73_USA SMH2013 accession) were germinated and grown under long days conditions (16:8 light:dark) at 25°C:20°C (light:dark) during seven days and harvested 4–6 h after Zeitgeber Time 0 (ZT0, lights on). The 2nd leaf containing inner leaves was flash frozen in liquid nitrogen, and stored at −80°C until use.

### Adenine methylation of gDNA using EcoGII

Methylation of 1µg of gDNA was performed at 37°C during 1h in the presence of 0.1 units/µl of EcoGII, in rCutSmart buffer in presence of 160 µM SAM, followed by heat inactivation at 65°C during 10 minutes. Methylation efficiency was measured by qPCR after DNA digestion with the m6A-specific restriction enzyme DpnI. 1µg of DNA was used to prepare ONT libraries using LSK110 kit following manufacture indications. Library was loaded in a FLO-MIN106D and runned in a MK1C device during 72hs.

### SAM-seq protocol

For Arabidopsis SAM-seq experiments, 250mg of seedlings were ground under liquid nitrogen and resuspended in 25 ml of extraction buffer 1 (sucrose 0.4M, Tris-HCl pH8 10mM, MgCl2 10mM, ethidium bromide 5mM). Tissues are fixed with 1% formaldehyde for 5 minutes in ice with gentle mixing, followed by the addition of 1.25 M glycine to twice the molar concentration of formaldehyde, materials sit in ice for 5 minutes before centrifugation at 4000 g for 20 minutes at 4°C. For nuclear isolation the pellet was resuspended in 1 ml Extraction Buffer 2 (sucrose 0.25M, Tris-HCl pH8 10mM, MgCl2 10mM, ethidium bromide 5mM, Triton x-100 1%, protease inhibitor 1X) and centrifuged at 4 °C for 10 min at 11000*g*. The pellet was then resuspended in 0.3ml Extraction Buffer 3 (sucrose 1.7M, Tris-HCl pH8 10mM, MgCl2 2mM, ethidium bromide 5mM, Triton x-100 0.15%, protease inhibitor 1X), transferred to a clean tube containing 0.3ml Extraction Buffer 3 and centrifuged at 4 °C for 30 minutes at 11,000*g*. To optimise MTase activity, two consecutive washes with 1ml of nuclei preparation buffer (MOPS pH7 20mM,NaCl 40mM, KCl 90mM, EDTA 2mM, EGTA 0.5mM, spermidine 0.5mM, spermine 0.2 mM, protease inhibitor 1X) and 1 ml of plant tween wash buffer (Tween-20 0.2%, NaCl 150mM, HEPES-KOH 20mM, spermidine 0.5mM, protease inhibitor 1X) were performed. The pellet of nuclei was resuspended in 200µl of activation buffer (1X rCutSmart, 800 µM SAM) containing 7.5U of EcoGII MTase. Samples were incubated under rotation at 37°C for 30 minutes. We noted that 30 minutes incubation with 37.5 U/ml of EcoGII provides the best signal/noise ratio to chart accessibility at ACRs and nucleosome profiling. For reverse cross-linking, 1 M NaCl was added to 200µl of 1X cutSmart to reach a salt concentration of 200mM. Samples were incubated at 65 °C for 1 hour for lysis to reverse cross-linking. 0.3mg/ml of RNAse and 0.1mg/ml Proteinase K were added to the nuclei and incubated for 40 minutes, followed by a phenol/chloroform DNA extraction.

For maize SAM-seq experiments, seven days old leaves were ground under liquid nitrogen and 750 mg of powder resuspended in 12.5ml of extraction buffer 1. Nuclei purification, fixation, and m6A-MTases reaction was performed as described above with the following modifications. Fixed nuclei’s pellet was resuspended in 38 µl 1X rCutSmart, containing 10µl SAM and 6µl EcoGII (i.e. 7.5U for 100µl) and incubated for 30 minutes at 30°C. For reverse cross-linking, 20 µl NaCl (5M) were added followed by an overnight incubation at 65 °C. DNA extraction was performed as described above.

#### ONT library preparation and sequencing

For each sample, between 0.5 and 1 µg DNA was used for library preparation using the Ligation Sequencing Kit SQK-LSK109. The library was prepared following the manufacturer’s instructions with minor modifications. The end-repair incubation time was increased to 20 min. Ligation incubation times were increased to at least 1h. Elution was always performed in 10 mM Tris-HCl. LFB was used for the final size selection step. DNA yield was quantified using the Qubit dsDNA HS Assay Kit (Q33230). Sequencing was performed on MinION with R9.4.1 flow cells (FLO-MIN106.1), or on PromethION with R9.4.1 (FLO-PRO002) or R10 (FLO-PRO114M) flow cells (Supplementary Table 1).

#### Detection of m6A and 5mC

For R9.4.1 chemistry, m6A base calling and mapping were performed using Megalodon with the Rerio model res_dna_r941_min_modbases-all-context_v001.cfg and default parameters. Adenines with methylated probability greater than 0.75 were considered as m6A. Otherwise, adenines were regarded as unmethylated. For 5mC methylation calling, fast5 files were squiggled with the FASTQ files obtained from Megadolon using Tombo v1.5.1. 5mC methylation was called from the squiggled fast5 files with DeepSignalPlant v0.1.6 (13) using the model model.dp2.CNN.arabnrice2-1_120m_R9.4plus_tem.bn13_sn16.both_bilstm.epoch6.ckpt. Single-molecule cytosine sites with predicted scores greater than 0.5 were considered as methylated. For R10 chemistry, m6A and 5mC base calling and mapping were performed using Dorado with the model dna_r10.4.1_e8.2_400bps_hac@v4.3.0 and default parameters. Adenines with methylated probability greater than 0.75 were considered as m6A. Methylation levels of genomic positions were calculated using the ratio of m6A/(m6A + A), where A denotes adenines predicted as unmethylated. To evaluate the dependency between 5mC and m6A, we compared the m6A levels over 20bp window between windows with different 5mC levels (5mC/C). Only windows with at least 3 cytosines were considered.

#### m6A methylation likelihood normalisation

To elucidate the intrinsic sequence preferences of EcoGII, we scanned the Col-CEN assembly to identify all adenine positions from both DNA strands and retrieved the 6-base sequences around (12-mers). To obtain normalisation constants, identical 12-mer sequences, irrespective of their strand-of-origin, were grouped together. For each of these 12-mer sequences we quantified the percentage of methylation of the focal adenines based on the EcoGII-treated DNA experiment, yielding a m6A methylation likelihood for each 12-mer. These values were re-centered to range between 0 and 1, and used to correct m6A levels of Adenines in both Watson and Crick genomic strands based on their specific K-mers. Similar procedure was performed for the Maize genome.

#### Analysis of SAM-seq over nucleosomes

Nucleosome positioning was inferred using MNase libraries previously published (14–17). MNaseq datasets were aligned to the Col-CEN reference genome assembly (18) and libraries with high mappability efficiency and median fragment lengths distribution of ∼147 bp were retained. Paired-end reads were mapped to the Col-CEN assembly using Bowtie v2.2.5 (19) with the following parameters: -N 1 --end-to-end --no-mixed --no-discordant. The resulting BAM files were sorted with samtools 1.6, and then converted to bigwig files using deeptools v3.5.1 (20) --binSize 1 --normalizeUsing RPGC --extendReads --effectiveGenomeSize 131559676 --ignoreForNormalization chrC chrM. The obtained bigwig files were pooled together to yield a smoothing effect. Next, an in-house script was implemented to perform peak calling using functions offered by the Numpy and Scipy libraries, (21, 22). Briefly, this calling first applied a convolution along the pooled profile using a 1D Gaussian kernel (standard deviation=30), and then calculated the second derivative of the convolved profile to determine local maximas. The mid-point of these maximas were defined as nucleosome dyads. The K-mean algorithm (K=2) based on dynamic time warping distances (23) was employed to clusterized the called dyads into well-positioned and poorly-positioned. Concretely, clustering used as input signals encompassing the 1 kb convolved profile upstream and downstream of each called dyad. The well-positioned dyads were further classified based on their genomic localization into: genes, TEs and CEN180s. Dyad’s positions were used to anchor metaplots generated by averaging MNase as well as normalised SAM-seq accessibility, mCG, mCHG, and mCHH levels. For visualisation, SAM-Seq accessibility metaplots were protected onto the cryo-EM nucleosome structure 7by0 (24).

#### Accessibility heterogeneity

Accessibility heterogeneity: To assess the accessibility heterogeneity of chromatin states we calculated the entropy of a probability distribution of the average density of m6As along reads mapped to genomic regions belonging to a specific chromatin state. The average methylation levels of a region were represented as a histogram by categorizing them into four discrete bins: 0.00-0.25, 0.26-0.50, 0.51-0.75 and 0.76-1.00. Normalizing the histogram to sum up one, we obtained a probability distribution. We then computed the Shannon entropy of this distribution. This entropy involves the summation of the probability associated with each discrete bin multiplied by the logarithm of this probability, where a bin represents a specific range of 6mA methylation level. Strictly speaking, the Shannon entropy is defined as: 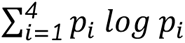, where *p_i_* denotes the probability associated with the *i*th bin. The resulting entropies for each chromatin state were depicted using violin plots

#### DNA footprints detection with SAM-seq

Deeptool was used to cluster ABI5 sites identified by DAP-seq based on SAM-Seq accessibility signal. A permutation test was conducted by allocating these sites across the clusters randomly. Per cluster enrichment of in vivo ABI5 binding [21] at DAP-seq sites, as well as the average ChIP-seq MACS2 score at these sites, was compared between the observed and random clusters. This procedure was repeated 500 times.

## RESULTS

### Simultaneous detection of m6A and 5mC in genomic DNA

Genome-wide simultaneous detection of m6A and 5mC using ONT sequencing has been successfully obtained in mammalian genomes (5, 9). To test whether this is also feasible in plants, we incubated Arabidopsis genomic DNA (gDNA) in a buffer containing the m6A-MTases EcoGII and the methyl donor S-adenosylmethionine (SAM) (Supplementary Figure 1), followed by ONT sequencing (Figure 1A). Genomic DNA (gDNA) without any treatment was also sequenced as control. We obtained long reads with a N50 of 5000 nt, with the longest reads reaching 200,000 nt. We processed these reads using two DNA modification callers, Megalodon (with Rerio model) and DeepSignalPlant (13), to detect 6mA and 5mC modifications, respectively (see Methods). We estimated 2% and 65% m6A methylation levels in control and EcoGII-treated gDNA, respectively, indicating low false positive calls and extensive EcoGII methylation. To test whether the presence of 5mC can affect EcoGII activity and/or m6A calling, we assessed m6A levels on 20-bp sequences extracted genome-wide having variable levels of 5mC methylation. Overall, we found little dependency between 5mC and m6A levels (Supplementary Figure 2).

**Figure 1.**
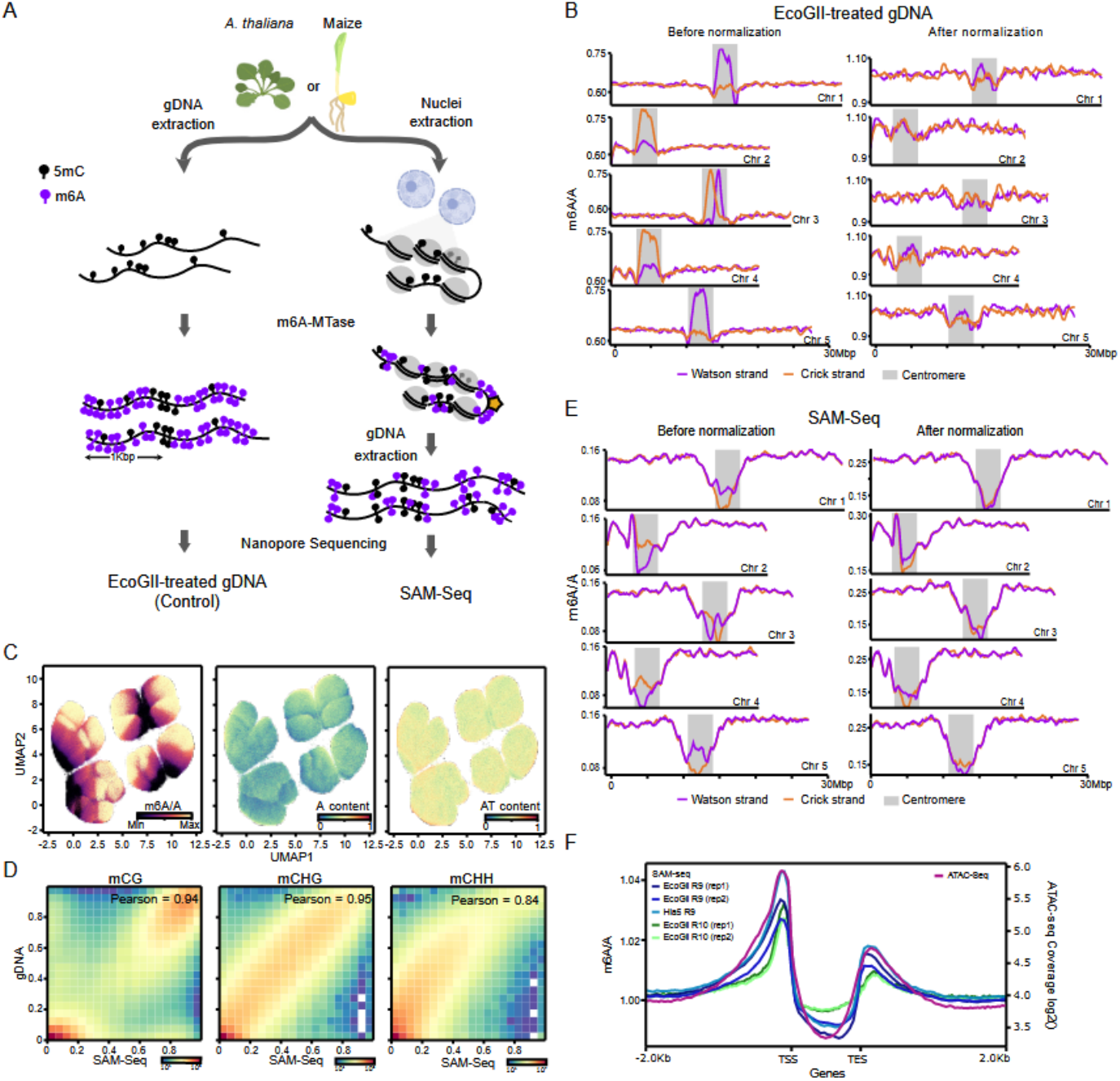
SAM-seq probes simultaneously chromatin accessibility and DNA methylation on plant tissues. **A**. Workflow of the SAM-seq protocol and assessment of m6A-MTase sequence preferences on gDNA (EcoGII-treated gDNA). **B**. Strand-specific m6A levels of EcoGII-treated genomic DNA before and after normalisation for m6A-MTase preferences over the five *A. thaliana* chromosomes. Centromeres are depicted as grey boxes. **C** UMAP projections of Arabidopsis 12-mer sequences coloured by m6A/A levels of EcoGII-treated gDNA, A content, and AT content. **D** Density plots and correlation analysis between ONT sequencing of gDNA and SAM-Seq for mCG, mCHG, and mCHH levels. **E**. Strand-specific SAM-seq m6A levels before and after normalisation for m6A-MTases preferences. **F**. Metaplot of Arabidopsis genes displaying ATAC-seq accessibility (Lu et al. 2017) (right Y axis) and SAM-seq chromatin accessibility (m6A/A) obtained from independent experiments using distinct m6A-MTases (EcoGII or Hia5) as well as distinct ONT chemistry (R9.4.1 or R10) (left Y axis).

We investigated the chromosome-wide profile of m6A levels of EcoGII-treated gDNA. Because no DNA-associated proteins are present in this *in vitro* experiment, m6A levels are expected to be uniform across the Arabidopsis genome. Nonetheless, we found uneven levels of m6A along chromosomes, together with strong strand-biased m6A methylation (Figure 1B). Strand-biased m6A levels were most notable at centromeres, which are made of highly repetitive centromeric satellites. To test whether these biases may be due to m6A-MTase sequence preferences, we modelled the m6A probability as a function of sequence composition, by calculating the average m6A level from EcoGII-treated gDNA across all potential 12-mer in the Arabidopsis genome. Dimensionality reduction analysis of the 12-mers space revealed several clusters of k-mers with high and low m6A levels, suggesting complex dependencies between nucleotide content and m6A (Figure 1C). Sequence biases were also evident over longer k-mers (Supplementary Figure 3). In addition, we observed strand-specific m6A levels in association with sequence content (Supplementary Figure 4). In combination, these results demonstrate the existence of significant m6A-MTase sequence preferences, likely underlying the uneven and strand-biased levels of m6A along chromosomes. Based on the 12-mers model, we generated a genome-wide map of m6A methylation likelihood as normalisation factor. After kmer-based normalisation, m6A levels were largely uniform along chromosomes (Figure 1B), indicating that the strand-specific m6A levels observed at centromeric repeats were largely driven by m6A-MTase sequence preference. In combination, our results demonstrate that m6A-MTase treatment combined with ONT sequencing provides a powerful method to detect simultaneously m6A and 5mC on long DNA molecules from plants, and underscore the importance of considering inherent biases in m6A-MTases activities to obtain reliable m6A profiles.

### Simultaneous chromatin accessibility and DNA methylation sequencing of Arabidopsis plants

To establish a robust protocol to probe chromatin accessibility in plant tissues, we purified, cross-linked, and permeabilized nuclei from two weeks old Arabidopsis seedlings (Figure 1A). m6A-MTases, such as EcoGII or Hia5, are known to be inhibited by detergents such as Triton or SDS (25) (Supplementary Figure 5), which are commonly used for plant nuclear extraction and permeabilization. In animal cells, these detergents can be replaced by digitonin, which does not affect m6A-MTases activity and interacts specifically with cholesterol (25). However, because cholesterol is present at very low concentrations in plant membranes (26), we tested the capacity of different combinations of soft detergents and washes to permeabilize Arabidopsis nuclei as well as their effect on m6A-MTases activity. Methyltransferase activity was tested by incubating gDNA extracted from treated chromatin with the restriction enzyme DpnI, which cuts GATC sites only in the presence of m6A (Supplementary Figure 1). We found that nuclei permeabilization with 1% Triton X-100 followed by two washing steps with 0.2% Tween-20 provides robust m6A methylation activity.

To investigate the performance of our SAM-seq approach to chart simultaneously chromatin accessibility and DNA methylation genome-wide in plants, we performed ONT sequencing of gDNA extracted from EcoGII-treated nuclei. The 5mC levels measured on gDNA extracted from EcoGII treated nuclei were highly consistent with those obtained from gDNA (Pearson correlations ranging from 0.84 to 0.95, Figure 1D), demonstrating efficient DNA methylation profiling by SAM-seq in the three cytosine contexts. Visualisation of normalised m6A levels revealed two broad chromosome compartments with high and low m6A-MTase accessibility, corresponding respectively to chromosome arms and heterochromatin centromeric and pericentromeric regions (Figure 1E). Notably, normalised m6A profiles showed little strand-biased m6A levels over centromeric regions, compared to non-normalized landscapes, highlighting the importance of the normalisation strategy for reliable accessibility profiling.

We next explored how incubation time and concentration of m6A-MTases affect SAM-seq accessibility results. We found that increasing incubation times improves profiling resolution over well-positioned nucleosomes (Supplementary Figure 6), at the cost of decreasing selectivity towards accessible chromatin regions (ACRs) upstream of the transcriptional start site (TSS) of genes. These results are consistent with previous observations made in animal cells [6]. To assess the robustness of our SAM-seq protocol, we compared the accessibility profiles obtained by SAM-seq with that obtained by ATAC-seq based on short-read sequencing (12). Overall, we found good agreement between both techniques (Figure 1F and Supplementary Figure 7), indicating that SAM-seq is a reliable method to chart chromatin accessibility at genes. Furthermore, accessibility profiles around TSSs of genes mirrored the nucleosome density assessed by MNase-seq (14), suggesting that SAM-seq can be used to detect nucleosome depleted regions (Supplementary Figure 8). We next explored SAM-seq reproducibility across independent experiments using EcoGII, sequenced either with the old (R9.4.1) or new (R10) nanopore chemistry, or performed with another m6A-MTase, Hia5 (Figure 1F). Last, we assessed directly the reproducibility of SAM-seq by performing pairwise comparisons between biological replicates. These analyses show good agreement among replicates, which, as expected, increased in relation to the depth of sequencing. An average vertical coverage of at least 17 informative adenines per strand leads to correlation of 0.93 (Supplementary Figure 9), demonstrating the high reproducibility of SAM-seq experiments.

### Accessibility and DNA methylation landscapes of chromatin states

To test whether SAM-seq can provide information about gene activity, we assessed normalised m6A levels around TSS of genes with contrasting expression levels. Accessibility increases in relation to gene expression, with lowly and highly expressed genes showing the lowest and highest m6A-MTase accessibility, respectively (Supplementary Figure 10). These results indicate that SAM-seq-based chromatin accessibility is a reliable proxy for gene activity. To explore further the ability of SAM-seq to chart chromatin landscapes, we plot m6A-MTases accessibility and DNA methylation over the nine chromatin states (CS) previously identified in Arabidopsis based on 16 genomic and chromatin features (27) (Figure 2A). These CSs distinguish promoters (CS1 and 2), upstream regulatory sequences (CS4), active (CS3) and inactive genes (CS6 and 7), intergenic sequences (CS5), and heterochromatic sequences (CS8 and 9). As expected, SAM-seq accessibility was high at active promoters (CS1), while the body of inactive genes and TEs have the lowest accessibility (CS7 and CS9) (Figure 2A, B and Supplementary Figure 11). Overall, chromatin accessibility correlates negatively with DNA methylation, consistent with a role for this modification in limiting chromatin accessibility (28). Nonetheless, CS8, which is enriched on TEs and characterised by the presence of DNA methylation and the PRC2-dependent facultative heterochromatin mark H3K27me3 (27), shows some level of accessibility. Interestingly, unlike CS9, which display the lowest accessibility and highest 5mC levels in the three sequence contexts, CS8 has high mCHH but reduced mCG and mCHG (Figure 2A), suggesting that the latter may be targets of the siRNA-directed DNA methylation pathway (RdDM). Simultaneous targeting of these *loci* by RdDM and PRC2 is consistent with recent studies demonstrating *in vivo* interaction of MORPHEUS’ MOLECULE1 (MOM1) and AIPP3 factors (29, 30), which are respectively involved in RdDM-mediated methylation and H3K27me3 recognition. In combination, these results provide evidence that CHH methylation, in combination with H3K27me3, might constitute a form of relaxed heterochromatin.

**Figure 2.**
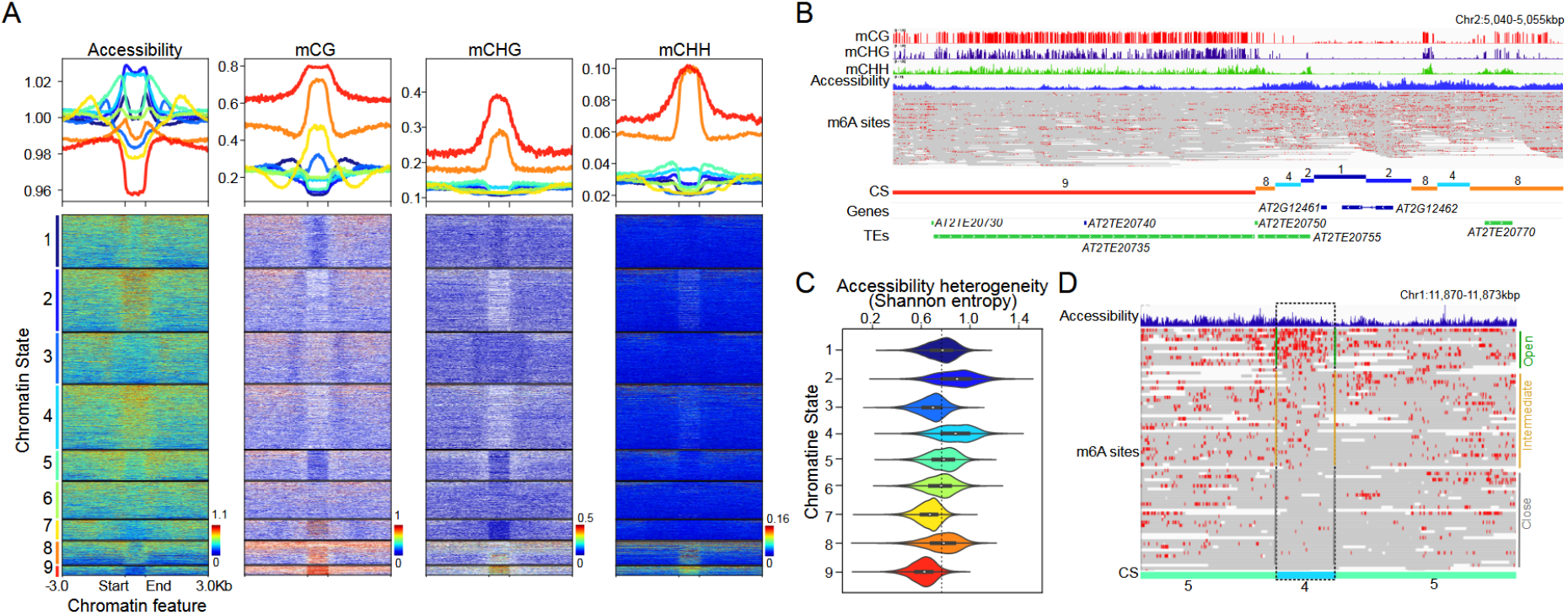
Accessibility and DNA methylation landscapes of Arabidopsis chromatin states. **A**. Accessibility, mCG, mCHG, and mCHH levels across nine chromatin states (Sequeira-Mendes *et al*., 2014). **B**. Genome browser view of SAM-Seq accessibility, mCG, mCHG, and mCHH levels. m6A along reads are indicated in red. **C**. Distribution of SAM-seq accessibility heterogeneity (Shannon entropy) across reads between CS locus. **D**. Genome browser view of SAM-Seq accessibility and m6A sites along reads spanning a CS4 locus with high accessibility entropy. Reads showing high (open), intermediate, and low (close) m6A levels are indicated.

To gain insights into the association between chromatin states and cellular heterogeneity, we took advantage of the ability of SAM-seq to probe accessibility across single molecules (Figure 2B). We found that broadly inactive genes (CS7) and silenced TEs (CS9) show the lowest levels of intermolecular m6A heterogeneity (Figure 2C and Supplementary Figure 12), indicating homogeneous chromatin accessibility across cells. Conversely, and despite their overall high accessibility, CS2 and to some extent CS4, which are characterised by bivalent marking of the antagonistic histone modifications H3K4me2 and H3K27me3 (27), display the highest 6mA entropy, with some loci showing on/off accessibility patterns across individual fibres (Figure 2C and 2D). The higher accessibility entropy at these loci provides strong evidence that co-marking reflects extensive cellular heterogeneity, rather than uniform chromatin states across fibres. Hence, CS2 and CS4 more likely reflect cellular-specific epigenetic regulations. Altogether, these results illustrate the power of SAM-seq to investigate epigenetic regulations within and across molecules, genomewide.

### Uncovering DNA footprints at cis-regulatory elements

Accessibility assays can be used to uncover DNA footprints (31), sequences that are locally protected by the binding of chromatin associated proteins such as transcription factors (TFs). Due to the high frequency of adenine-thymine base pairs in the Arabidopsis genomes (about 64%), the high resolution provided by m6A-MTase is ideal to assess short-scale changes in accessibility, such as those associated with DNA footprints. To determine whether m6A-MTase can outline TF-binding sites (TFBS), we explored SAM-seq profiles around 390 putative TFBSs identified by *in vitro* assays (32). Overall, we detected higher SAM-seq accessibility in the vicinity of 90 of these TFBS motifs (Supplementary Figure 13), consistent with numerous functional regulatory sequences residing within active chromatin regions. Furthermore, we found that SAM-seq accessibility drops at TFBSs were evident for at least 40% of these TFs, suggesting the presence of DNA footprints (Supplementary Figure 13).

To evaluate the potential of SAM-seq to pinpoint occupied TF binding sites *in vivo*, we focused on ABA INSENSITIVE 5 (ABI5), which is a basic leucine zipper (bZIP)-type TF involved in the post-germination response to abscisic acid. Clustering of predicted ABI5 binding sites based on SAM-seq accessibility yielded three types of sites with strong, intermediate, or no protection footprint (Figure 3A). To test the significance of the clustering, we analysed *in vivo* occupancy of ABI5 determined by ChIP-seq (32). Sites with strong and low accessibility protection were highly and lowly overrepresented with ABI5 ChIP-seq peaks, respectively (Figure 3B). Furthermore, the intensities of ChIP-seq peaks were much higher at sites with accessibility footprints than those lacking such footprints (Figure 3C). Next, we assessed the minimum ONT sequencing depth required to detect accessibility footprints. To this end we subsampled SAM-seq sequencing data, clustered ABI5 sites based on normalised m6A levels, and quantified ChIP-seq peaks enrichment and intensities as performed above (Supplementary Figure 14). This analysis established that 1.25Gbp of ONT data (about 10X genome-wide coverage) is sufficient to robustly detect ABI5 footprints in Arabidopsis. Altogether, these results demonstrate the power and sensitivity of SAM-seq to identify occupied TFBS sites *in vivo*.

**Figure 3.**
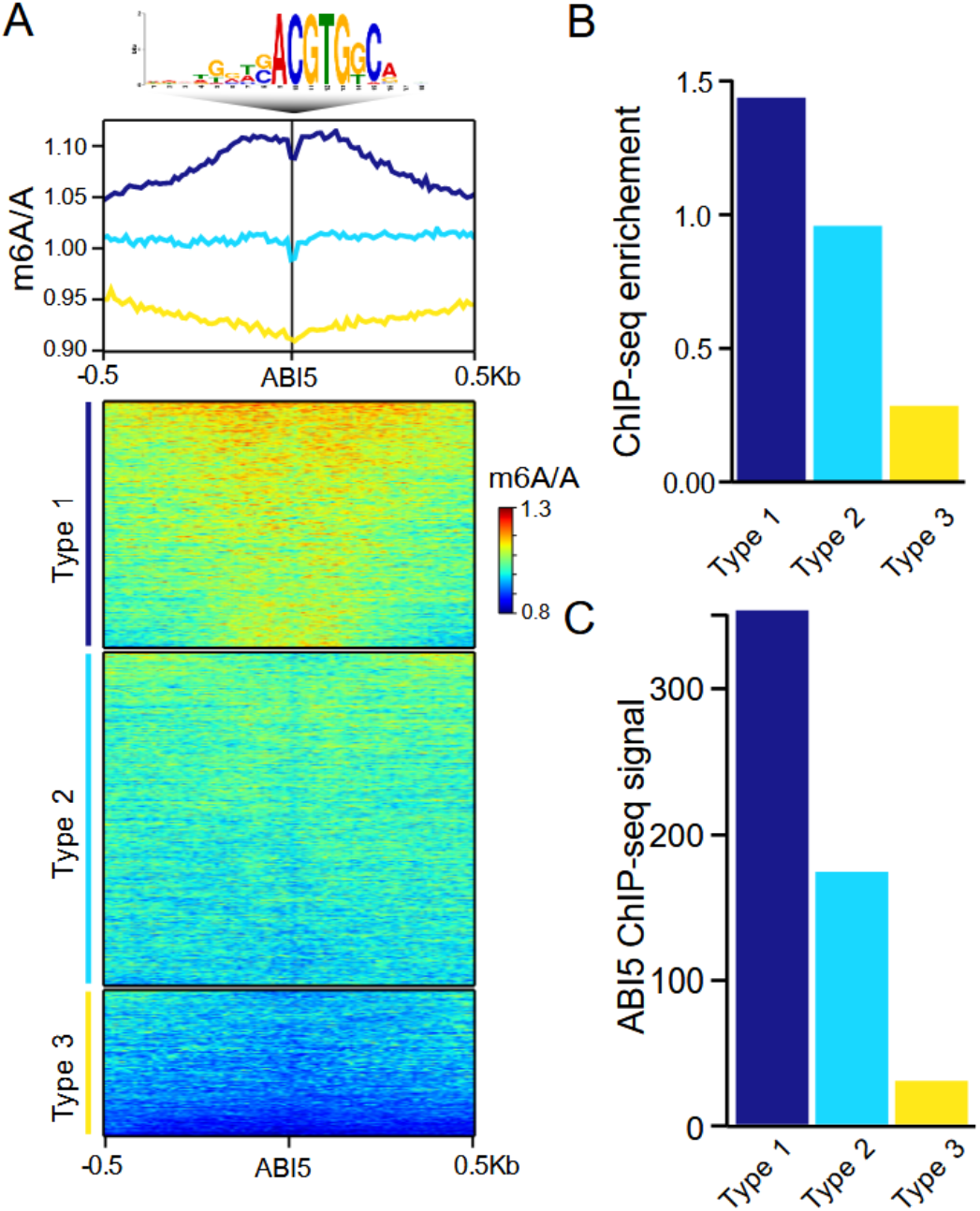
SAM-seq footprinting pinpoints sites bound by TFs *in vivo*. **A**. Metaplot and heatmaps of SAM-Seq chromatin accessibility around putative ABI5 binding site determined in-vitro by DAP-seq. Clustering of chromatin accessibility profiles identified three types of ABI5 bindings sites. **B**. Enrichments of ABI5 Chip-Seq peaks at type 1, 2 or 3 transcription factor binding sites (TFBS). **C**. ABI5 Chip-Seq peaks signal over type 1, 2 or 3 TFBS.

### Fine-grained accessibility and DNA methylation of nucleosomes

Accessibility assays have the potential to provide information on nucleosomal positioning. To test whether SAM-seq can provide such information in plants, we analysed normalised m6A levels around well positioned nucleosomes (see methods). Because accessibility over nucleosomes may differ between euchromatin and heterochromatin, we distinguished between well-positioned nucleosomes at genes, TEs, and centromeric repeats. We observed the lowest m6A levels within core nucleosomal regions, with the notable exception of the mid-points of centromeric nucleosomes (Figure 4A). Accessibility peaks at the linker DNA situated approximately 75 base pairs upstream and downstream of the nucleosomal centres (*i.e.* dyads). Nucleosomal-scale SAM-seq protection extended several hundred base pairs in both directions from the focal nucleosomal midpoint, mirroring the MNase nucleosomal signal. The significant SAM-seq signal that we observed at the dyad of centromeric repeats (Figure 4B) suggests that CENH3-containing nucleosomes have unique accessibility patterns. Structural and biochemical studies suggested that DNA at the entry/exit sites of centromeric nucleosomes is highly flexible and does not cross above the dyad, thereby impeding the binding of linker histone H1 (33, 34). To further test this possibility, we compared SAM-seq accessibility over centromeric repeats with high and low CENH3 levels (18) and observed that accessibility at dyads was higher at CENH3-rich than CENH3-poor nucleosomes (Supplementary Figure 15). Altogether, our SAM-seq results suggest that flexible DNA ends of centromeric nucleosomes provide higher accessibility at dyads, with potential implications for the epigenetic regulation of centromeres.

**Figure 4.**
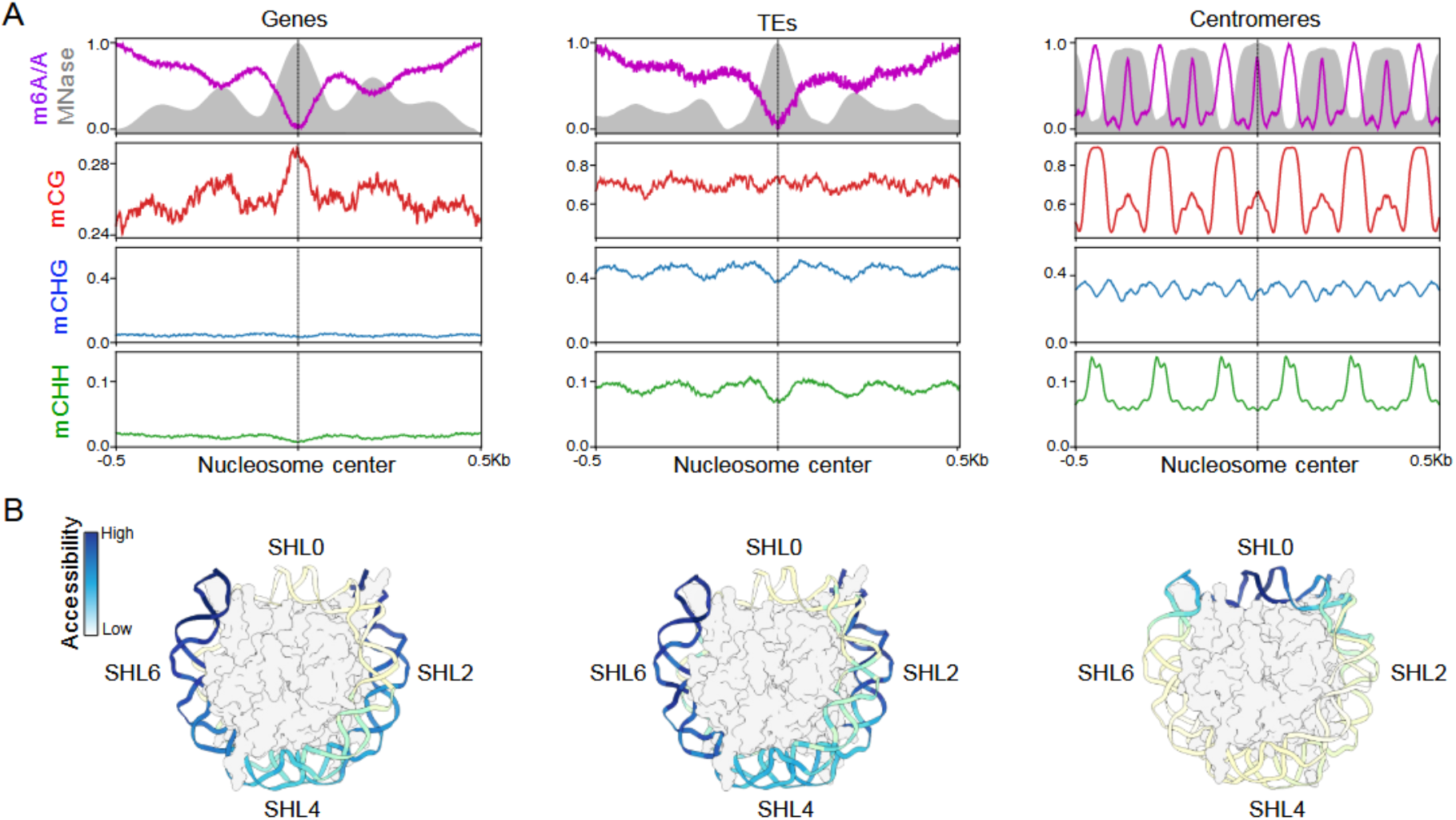
Fine-grained accessibility and DNA methylation of Arabidopsis nucleosomes. **A**. Metaplots showing MNase (grey area), SAM-Seq accessibility (purple line), mCG (red), mCHG (blue) and mCHH (green) over well-positioned nucleosomes in genes, TEs, or centromeres. **B**. Projection of SAM-Seq accessibility metaplots onto the cryo-EM nucleosome structure (7by0). Superhelical locations (SLH) 0, 2, 4, and 6 are indicated.

We next investigated how nucleosome accessibility affects DNA methylation. Over genes, nucleosomal DNA has higher levels of CG methylation compared to linker DNA (Figure 4A). This result is consistent with previous observations (14), and suggests that MET1 activity is not impaired by euchromatic nucleosomes. Conversely, linker DNA at TEs and centromeres show the highest methylation levels compared to nucleosomal DNA, supporting the notion that heterochromatin nucleosomes are strong barriers to DNA methyltransferases (14). In addition, methylation profiles also reveal that dyads of centromeric nucleosomes have high CG methylation, and that this methylation follows the nucleosomal accessibility landscape (Figure 4B). This result suggests that relaxed DNA wrapping of centromeric nucleosomes promotes the accessibility of MET1 at dyads, leading to high levels of CG methylation. The unusual accessibility and DNA methylation landscapes of centromeric nucleosomes revealed here may have important implications for centromeric function and chromosomes segregation.

### Chromatin profiling of Arabidopsis repeated sequences

Because long-reads can be unambiguously aligned over highly repetitive sequences, we explored the ability of SAM-seq to simultaneously chart chromatin accessibility and DNA methylation over such sequences, including *CEN180* centromeric satellites. Consistent with their role in kinetochore assembly, centromeric repeats show the lowest accessibility compared to euchromatic regions (Figure 1E). Recent targeted m6A-MTase-based profiling of centromeres reported intriguing accessibility patterns of *CEN180* repeats, including higher accessibility of *CEN180* compared to surrounding *ATHILA5* retrotransposons, strand-biased accessibility, and reduced accessibility of linker DNA of centromeric nucleosomes (10). We reasoned that such patterns could be influenced, at least in part, by the sizable sequence preferences displayed by m6A-MTases (Figure 1). Indeed, strand-biased m6A levels of *CEN180* repeats are evident on EcoGII-treated (naked) gDNA, and largely disappeared after controlling for m6A-MTase preference (Figure 1C and Supplementary Figure 16). Lower SAM-seq signal of *ATHILA5* retrotransposons compared to surrounding *CEN180* repeats also vanished after controlling for m6A-MTase sequence preference (Supplementary Figure 17. Furthermore, our SAM-seq profiles revealed higher accessibility of linker DNA (Figure 4). These results underscore the importance of considering enzymatic preferences for chromatin profiling, and provide evidence that accessibility patterns of Arabidopsis centromeres previously reported (10) can be largely explained by technical artefacts.

We next explored the accessibility and DNA methylation across TEs in the Arabidopsis genome. Overall, SAM-seq accessibility and DNA methylation correlate negatively (Figure 5A and B), in line with previous studies showing a dampening role of DNA methylation in TE accessibility (28). To explore further this relationship, we clustered TE sequences based on accessibility profiles. This unsupervised model identified six clusters of TEs (Figure 5A). Most clusters, except C1 and 3, contained lowly accessible TEs surrounded by regions with variable degrees of accessibility. As expected, the less accessible cluster, C6, has the highest levels of DNA methylation in all sequence contexts, is highly enriched on TEs belonging to *Ty3/Gypsy* families (Figure 5C), and are located far from genes (Figure 5D). Clusters 1 and 3, in contrast, have high accessibility and reduced DNA methylation levels. TEs in these clusters belong to Helitrons, SINEs, LINEs, and Pogo, and locate nearby genes, suggesting that they might serve as regulatory sequences. Potentially regulatory TEs have been recently identified based on dozens of chromatin marks (35). Comparison between our clusters of TEs defined solely on SAM-seq results with those obtained previously (35) confirmed that C1 and C3 largely span TEs enriched in chromatin marks associated with regulatory functions as well as TFBS (Supplementary Figure 18). Moreover, accessibility over repetitive sequences have been shown to facilitate meiotic double strand breaks generated by SPO11 topoisomerase-like complexes, with nucleosome-depleted Helitron/Pogo/Tc1/Mariner DNA transposons showing an enrichment in SPO11-1 oligos (36). As these types of TEs appear enriched in C1 and C3, we tested the accumulation of SPO11-1 oligos across the six TE clusters based on SAM-seq accessibility (Supplementary Figure 19). This analysis confirmed the strong association between TE accessibility and SPO11-1-oligos levels, and indicate that TEs in C1 and C3, which are enriched in chromatin marks associated with regulatory functions as well as TFBS, are hotspots of meiotic DSB. In combination, these results demonstrate that SAM-seq can be used to characterise chromatin states at repetitive sequences and to pinpoint TE-derived sequences with potential regulatory functions.

**Figure 5.**
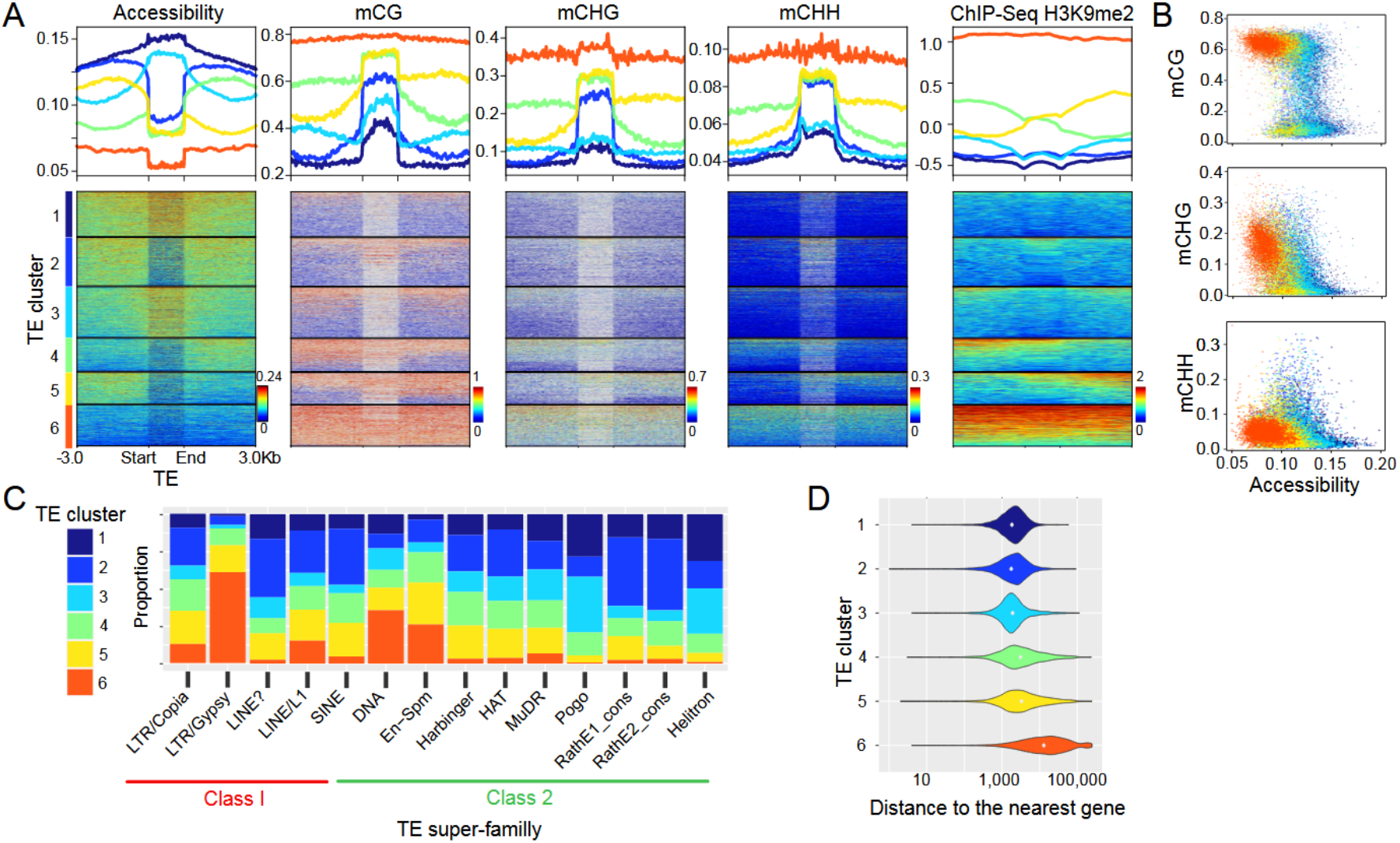
Chromatin profiling of Arabidopsis TEs. **A**. Metaplots and heatmaps for SAM-Seq accessibility, mCG, mCHG and mCHH methylation, as well as H3K9me2 from ChIP-seq data, across TEs in the Arabidopsis reference genomes. Clustering of TEs based on SAM-Seq accessibility identified six clusters. **B**. Dotplots showing SAM-Seq accessibility and mCG, mCHG or mCHH for each TE. TEs are coloured based on the clusters obtained in A. **C**. Proportion of TEs belonging to each cluster across TE superfamilies. **D**. Distribution of distances of TEs to the nearest gene across TE clusters.

### High-resolution chromatin profiling of complex genomes

With few exceptions, notably Arabidopsis, plant genomes carry a high load of repeats, making it very challenging to profile chromatin landscapes using short-read sequencing technologies. The long read-based approach employed by SAM-seq therefore provides an unprecedented opportunity to investigate the chromatin of large and complex plant genomes. We adapted SAM-seq to investigate chromatin accessibility and DNA methylation in the 2.1Gbp-long maize genome (see Methods), 80% of which corresponds to TE and other repeated sequences (37). We performed SAM-seq on maize leaves of the reference line B73 and sequenced using a PromethION R9.4.1 flow cell, yielding 52 million reads. Unambiguously mapped SAM-seq reads covered 96% of the reference maize genome, compared to only 50-64% typically obtained with short-read sequencing. The significant increase in horizontal coverage demonstrates the definitive advantage of using SAM-seq to investigate the epigenome landscapes of highly-repetitive plant genomes.

Overall m6A level was ∼3%, which is much lower than that obtained for Arabidopsis. This result is consistent with previous observations indicating that most of the maize genome is largely inaccessible (38). Notwithstanding the overall low accessibility, chromosome-wide normalised m6A levels were punctuated by narrow peaks of SAM-seq accessibility (Figure 6A and B). These peaks were often located near the TSS of genes (Figure 6B), and metaplots at genes recapitulated the patterns obtained by ATAC-seq and DNAseq (39, 40)(Figure 6A), consistent with these sites being ACRs. Accessibility profiles at genes TSS of another maize inbred line, M52, were similar to the ones obtained in B73 (Supplementary Figure 20), further supporting the reproducibility of SAM-seq to investigate maize chromatin. We then took advantage of the large maize genome, where ACRs can be located away from genes, to further test the correspondence between SAM-seq and ATAC-seq in B73. For this, we evaluated normalised m6A levels at high-confidence proximal (<2kb from TSS) and distal (>2kb from TSS) ACRs previously defined by ATAC-seq (39). This analysis showed that SAM-seq accessibility was the highest over ACRs, independently of their distance to genes (Figure 6C), thus highlighting that this method can inform on chromatin accessibility outside of genes, a feature particularly important for large plant genomes (41). In maize, ACRs have been found to be enriched in TFBSs (39). Consistent with these observations, SAM-seq accessibility increases around binding sites identified by DAP-seq for 32 TFs (32, 42) (Supplementary Figure 21). Last, we explored the SAM-seq accessibility of genes displaying different levels of expression (43) (Figure 6D). This analysis revealed a good correlation between SAM-seq accessibility at TSS of genes and expression levels. Altogether, these results demonstrate that SAM-seq is scalable to large genomes, enables the identification of ACRs at promoters, proximal and distal regions, and provides a reliable proxy for gene activity *in vivo* for complex plant genomes.

**Figure 6.**
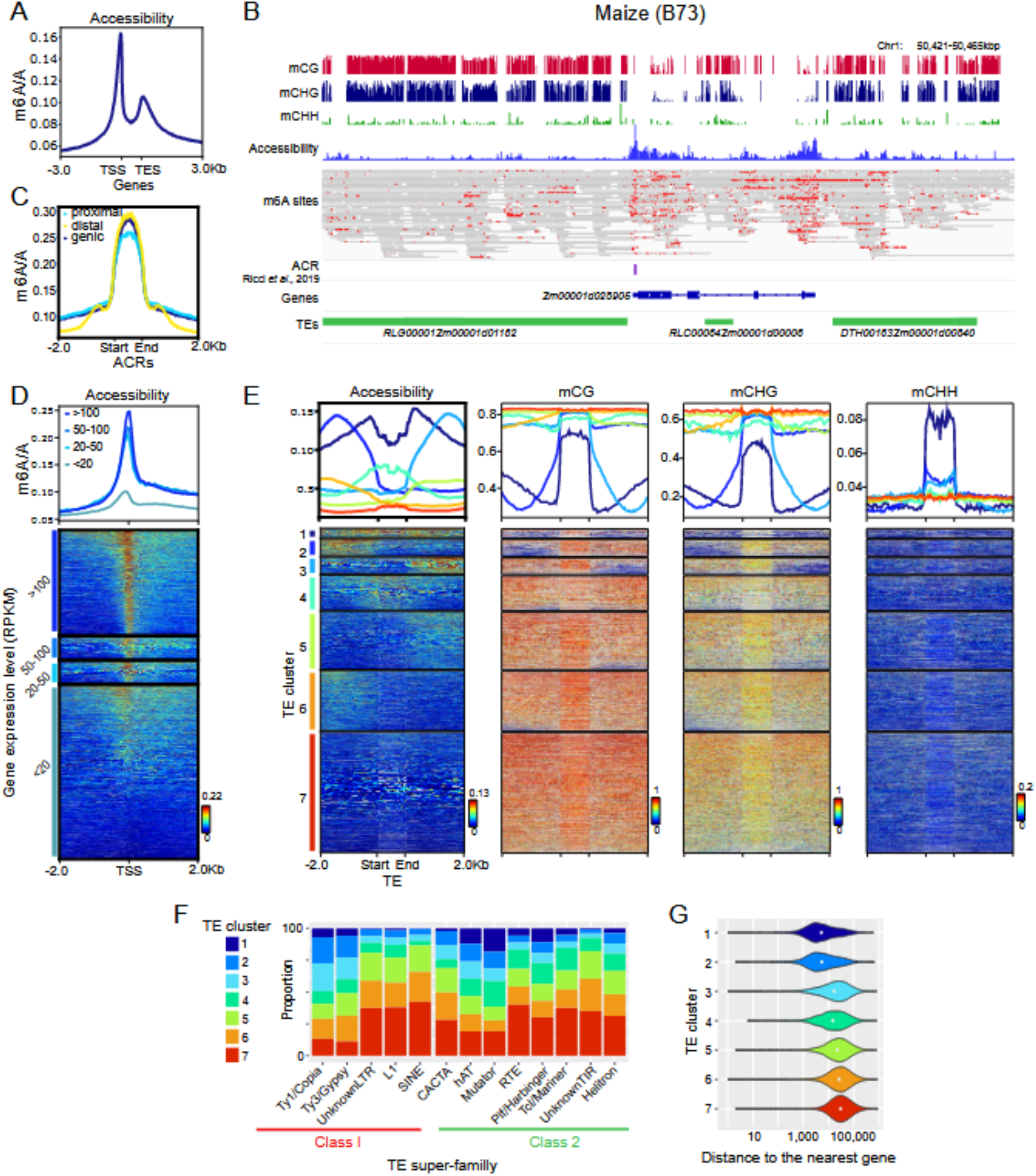
SAM-seq chromatin profiling of the maize genome. **A.** Metaplots of chromatin accessibility signal over genes. **B**. Genome browser view of SAM-Seq accessibility, mCG, mCHG, and mCHH levels. m6A along reads is indicated in red and accessible chromatin region (ACR) detected by ATAC-seq from Ricci *et al.,* 2019 are shown. **C.** Metaplots of chromatin accessibility signal over genic, proximal or distal ACRs. **D.** Metaplots and heatmap of chromatin accessibility signal over TSS with different expression levels. **E**. Metaplots and heatmaps for SAM-Seq accessibility, mCG, mCHG and mCHH methylation, across TEs in the reference Maize genomes. Clustering of TEs based on SAM-Seq accessibility identified seven clusters. **F**. Proportion of TEs belonging to each cluster across TE superfamilies. **G**. Distribution of distances between TEs and their nearest gene across TE clusters.

Given the high TE-content of the maize genome, we tested the ability of SAM-seq to chart accessibility and DNA methylation over TEs. Overall, we found that maize TEs are poorly accessible and associated with high mCG and mCHG levels, supporting previous observations based on limited genome coverage obtained through short-read sequencing technologies (44). Clustering of TE sequences based on SAM-seq chromatin accessibility identified seven clusters (Figure 6E). The largest cluster (C7) gathers TEs with, and embedded in regions of low accessibility, high mCG and mCHG, and low mCHH levels. These types of inaccessible, highly methylated TEs, are typically located far away from genes and associated with Harbingers, RTE, Helitrons and unknown LTR and TIR superfamilies (Figure 6F). Conversely, upstream and downstream regions respectively flanking TEs in C2 and 3, and to a lower extent C5 and 6, show relatively high SAM-seq accessibility and low CG and CHG methylation. Notably, these TEs have high mCHH levels, which peaked towards TE’s border facing the accessible flankings. Such a pattern is reminiscent of “CHH islands”, which has been described as narrow clusters of mCHH typically located nearby genes and proposed to operate as epigenetic boundaries between euchromatin and heterochromatin in maize (45). Last, Cluster 1, and to a lower extent C4 too, gathers regions with high chromatin accessibility both within TEs and at flanking regions (Figure 6E). TEs from this cluster have particularly high mCHH, reduced mCG and mCHG levels (Figure 6E), typically belong to *hAT*, *Mutator*, and *PIF/Harbinger* superfamilies, and are located close to genes (Figures 6F and G). Altogether, these results indicate that maize TEs preferentially methylated in the CHH context are not linked to reduced accessibility nor to euchromatin/heterochromatin boundaries. Conversely, these TEs may be playing a role as *cis*-regulatory elements of nearby genes, as has been recently proposed for the emergence of distal regulatory sequences across angiosperm species (41).

## DISCUSSION

The intricate interplay between nucleosome positioning, DNA methylation, and chromatin accessibility is fundamental for genome regulation in eukaryotic organisms. However, addressing these aspects has been so far challenging due to the technological limitations imposed by the use of short-read sequencing methods. This challenge is particularly critical when studying highly repetitive sequences, which are pervasive components of most plant genomes. Here, we adapted a method that takes advantage of m6A-MTases and long-read sequencing to ‘stencil’ accessible DNA (9) to simultaneously map DNA methylation and chromatin accessibility in complete plant genomes. We demonstrate that SAM-seq can be applied to obtain fine-grained chromatin landscapes in plant species with widely variable genome sizes and repetitiveness. Moreover, SAM-seq’s technical simplicity, reproducibility, and cost-effectiveness make it an ideal approach for investigating chromatin marks and their dynamics in plants with large genomes.

The dynamic range and power of SAM-seq are comparable between Arabidopsis and maize, illustrating its versatility across species and genome complexities. This versatility would enable the characterization of chromatin landscapes in virtually any plant species, providing unprecedented insights into chromatin regulation, gene activity, and nucleosome positioning. Furthermore, given the recent improvements in the quality and quantity of ONT sequencing outputs, SAM-seq offers an all-in-one tool to generate *de novo* genome assemblies together with detailed epigenome information. This will significantly facilitate the investigation of chromatin-based regulations across the tree of life, bridging the gap between model and non-model species.

One significant limitation in investigating chromatin accessibility has been the inherent sequence biases of DNA modifying enzymes, such as those previously observed for DNaseI and Tn5 retrotransposons (11, 12). Here, by subjecting genomic (naked) DNA to m6A-MTases, we uncovered significant sequence preferences. We also show that these preferences can generate spurious accessibility landscapes. For instance, m6A-MTase sequence preferences can lead to significant strand-specific accessibility profiles, especially over highly repetitive sequences. Previous work using EcoGII m6A-MTase reported only modest differences in methylation levels among k-mers (9). These prior findings may have resulted from an incomplete characterization of m6A-MTase preferences. Indeed, the significance of biases increases with K-mers size (9), and we uncovered strong biases when considering 12-mers and 20-mers. Hence, m6A-MTase activity might be likely influenced by regional DNA composition, rather than local sequence motifs. Given the rapid growth in the number of methods using m6A-MTases to investigate chromatin, including recent developments of antibody-driven targeting of m6A-MTases to map DNA binding proteins (25, 46), we underscore the importance of measuring and correcting inherent biases introduced by m6A-MTases preferences.

Chromatin accessibility is commonly assessed using ATAC-seq, which is based on short-read sequencing. However, detecting DNA footprints and nucleosome positioning using this technique remains challenging. We demonstrate that SAM-seq is a multipurpose chromatin profiling technique for plants capable of capturing a wide spectrum of chromatin nuances, from subtle, small-scale alterations to large-scale changes. This wide-resolution profiling enabled us to uncover striking variations in accessibility and DNA methylation along the nucleosome surface. In particular, we show that dyads of Arabidopsis centromeric nucleosomes are highly accessible, potentially facilitating the activity of the CG-specific DNA methyltransferase MET1. This atypical accessibility and DNA methylation nucleosome landscape may have important implications in the epigenetic regulation of centromeres and chromosomal segregation. In addition to mapping nucleosomes, the high-resolution of SAM-seq enables the detection of TFBS occupied in vivo. We found that at least one third of putative binding sites for the Arabidopsis TF ABI5 reside within open chromatin and show DNA footprints (Figure 4). Similar picture was obtained in maize, indicating that SAM-seq can be used to investigate cis-regulatory regions in different plant species.

The unique ability of SAM-seq to probe accessibility across single chromatin fibres open the opportunities to test for potential intermolecular variability, which should reflect variability at the cellular level. By comparing the intermolecular m6A entropy between chromatin states, we found that domains carrying antagonistic chromatin marks, the so-called bivalent states, are characterised by high heterogeneity of accessibility. Given that our SAM-seq experiments were performed using above-ground Arabidopsis tissues, the high heterogeneity we observed likely reflects tissue specific regulations of gene activity. Based on our findings, a key priority for the future will be to perform SAM-seq in different tissues and organs to investigate cell type-specific chromatin heterogeneity.

Clustering of TEs based on SAM-seq accessibility allowed us to distinguish between highly heterochromatin TEs, TEs located in heterochromatin/euchromatin boundaries between, TEs embedded in euchromatin, and bonafide euchromatic TEs. Remarkably, unlike Arabidopsis where DNA methylation in the three cytosine contexts correlate negatively with accessibility, euchromatic (i.e. accessible) TEs in maize show the highest mCHH levels. Given that maize lacks the mCHH-specific DNA methyltransferase CMT2, our results provide strong evidence supporting that accessible TEs are the preferred targets of RdDM (40). Furthermore, we found that mCHH and accessibility is higher at the edge of these TEs, and this pattern is mirrored by mCG and mCHG. This result suggests that RdDM may establish a transition zone between heterochromatin and euchromatin, preventing the spreading of DNA methylation over genes as well as preventing accidental PolII read-through transcription from nearby genes. In combination, the high-resolution accessibility and DNA methylation landscapes of SAM-seq offers a powerful toolkit to study the interplay between nucleosomes, chromatin accessibility, and DNA methylation. Its technical simplicity and ability to simultaneously characterise multiple chromatin features, ranging from small-scale changes to chromosomal-scale patterns, enables the investigation of chromatin-based regulations in virtually any plant species.

## Availability of data and materials

The datasets generated and/or analysed during the current study are available in the European Nucleotide Archive (ENA) repository <u>PRJEB69301</u>. Publicly available Arabidopsis ATACseq, MNaseq, H3K9me2 ChIPseq, DAP-seq and ABI5 ChIPseq data were obtained from GEO bioprojects <u>GSE85203</u>, <u>GSE96994</u>, <u>GSE139460</u>, <u>GSE190317</u>, <u>GSE207391</u>, <u>GSE60143</u>. Publicly available Maize ATAC-seq and DAP-seq were obtained from GEO bioprojects <u>GSE60143</u> and <u>GSE120304</u>. Genome-wide tracks of SAM-seq accessibility, mCG, mCHG, and mCHH for Arabidopsis and Maize are available at DOI:10.1101/2023.11.15.567180 and DOI:10.1101/2023.11.15.567180, respectively. All data generated or analysed during this study are included in this published article. Computer code to calculate K-mer normalisation is available at https://github.com/aedera/m6anormalization. Detailed SAM-seq protocols for Arabidopsis and maize are publicly available at dx.doi.org/10.17504/protocols.io.8epv5x1xdg1b/v1 and dx.doi.org/10.17504/protocols.io.kqdg3x25zg25/v1, respectively.

## Author contributions

LQ conceived the project. BL developed and implemented SAM-seq, performed most bioinformatic analysis with additional inputs from AE, and interpreted the data. AE performed m6A methylation likelihood normalisation and nucleosome analysis. CV generated maize samples and discussed results. LQ interpreted the data and drafted the manuscript, with additional inputs from all the authors. All authors read and approved the paper.

## Acknowledgements

We thank members of the Quadrana group for discussions. LQ thanks Vincent Colot for insightful discussions. We thank Nicolas Altemose for sharing the Hia5-pA enzyme. We thank the GDR 3546 Mobil-ET for fostering interactions between CV and LQ.

## Supplementary Data

**Supplementary Figure 1.**
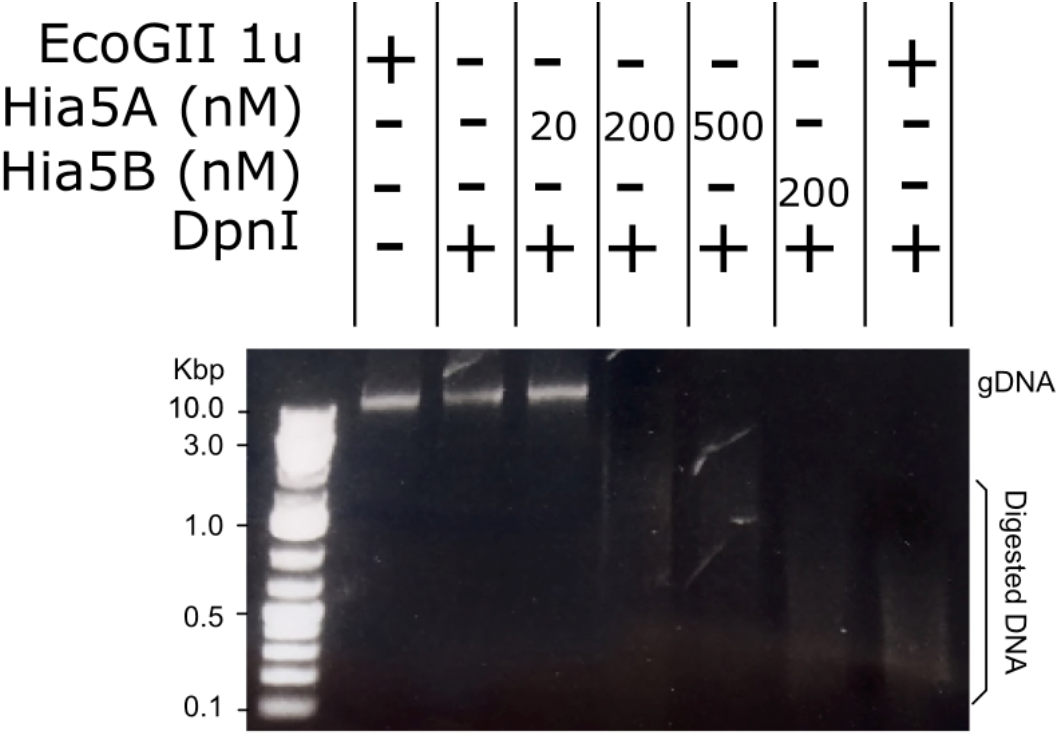
Assessment of EcoGII and Hia5 m6A-MTases *in vitro* activity on genomic DNA using the m6A-specific restriction enzyme DpnI.

**Supplementary Figure 2.**
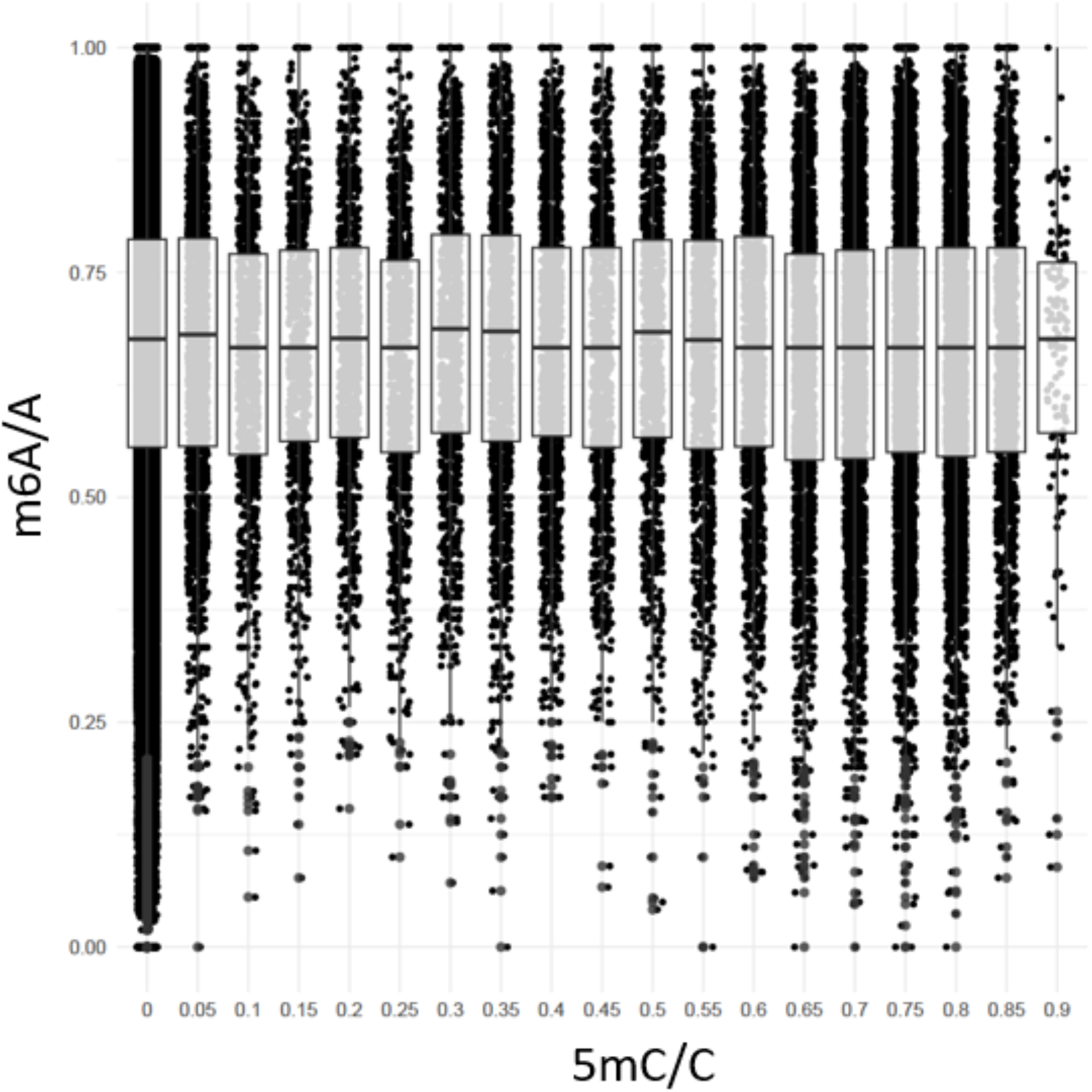
Scatter and boxplots displaying m6A methylation levels (m6A/A) per 20bp window with different 5mC levels (5mC/C) obtained on EcoGII-treated gDNA.

**Supplementary Figure 3.**
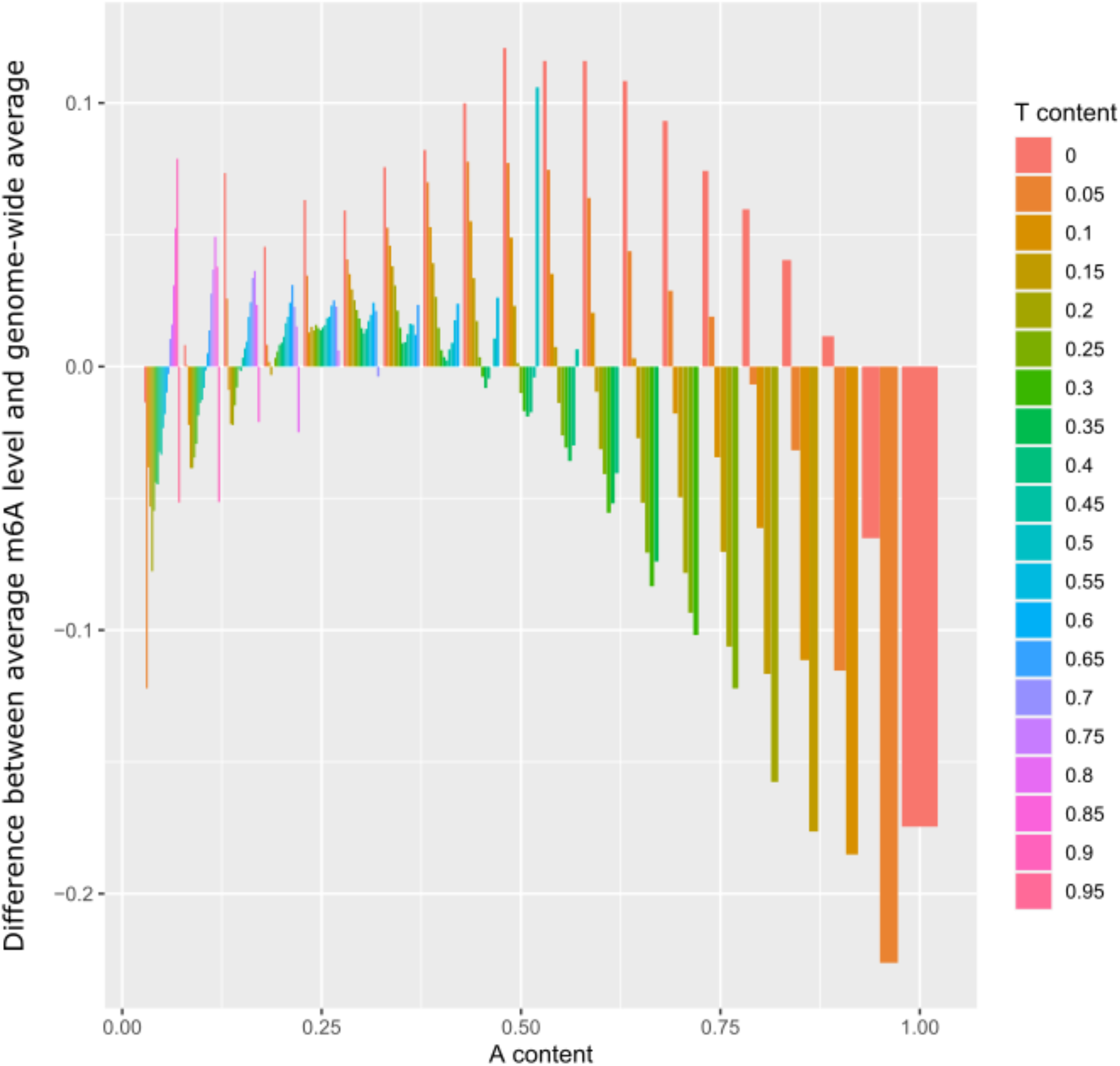
Deviation of average m6A levels of 20-mer in relation to A and T content. Differences between average m6A levels per k-Mer and the global average of m6A level are shown.

**Supplementary Figure 4.**
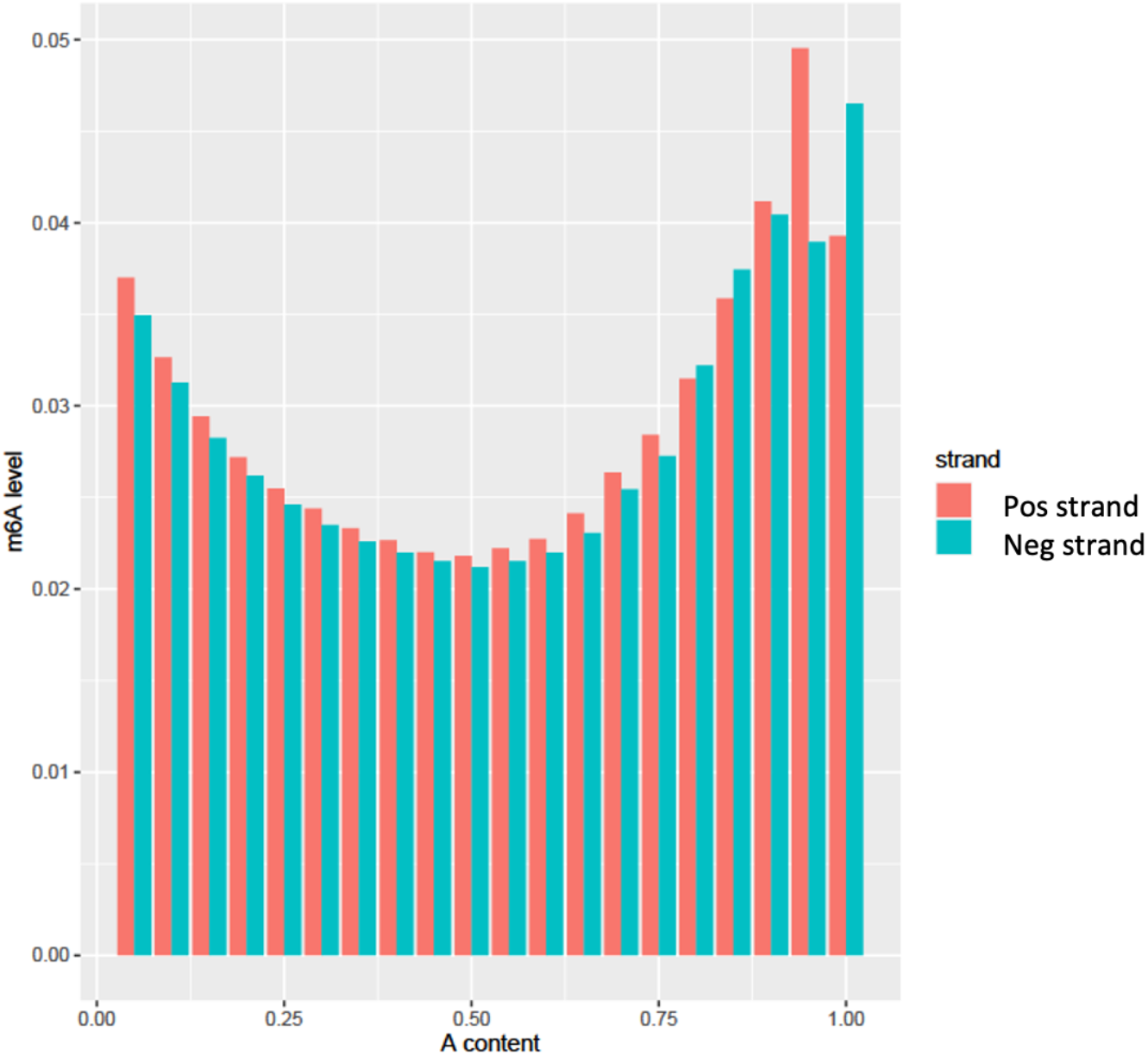
Strand-specific m6A levels over 20bp-long windows and in relation to A content of the window.

**Supplementary Figure 5.**
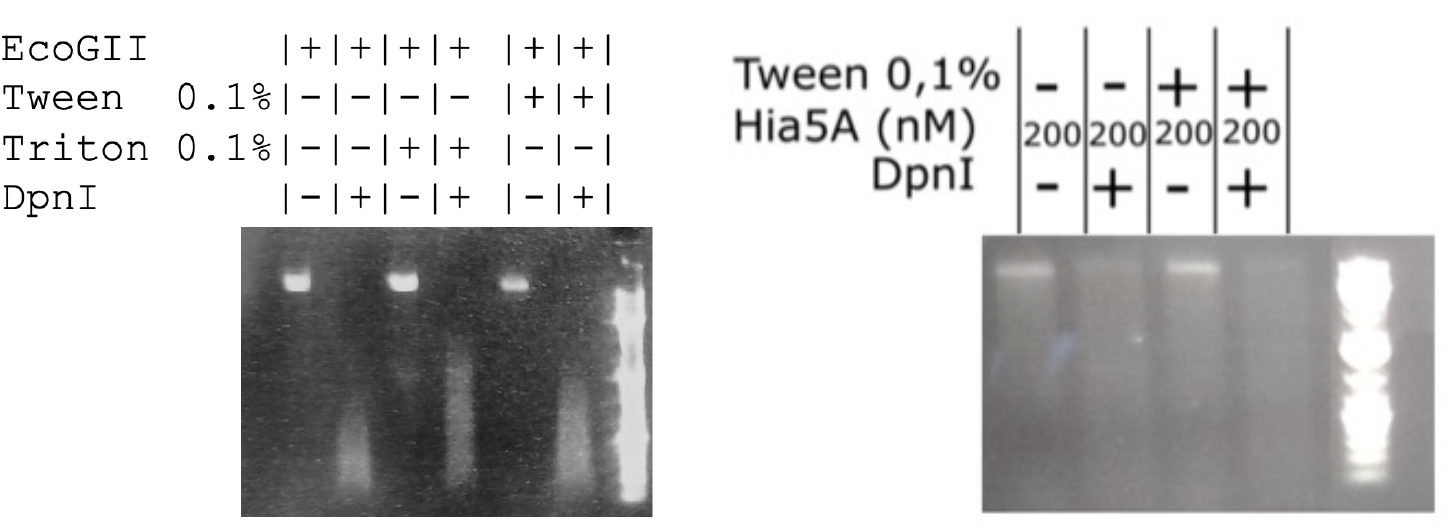
Testing the impact of different detergents on *in vitro* m6A-MTases activity.

**Supplementary Figure 6.**
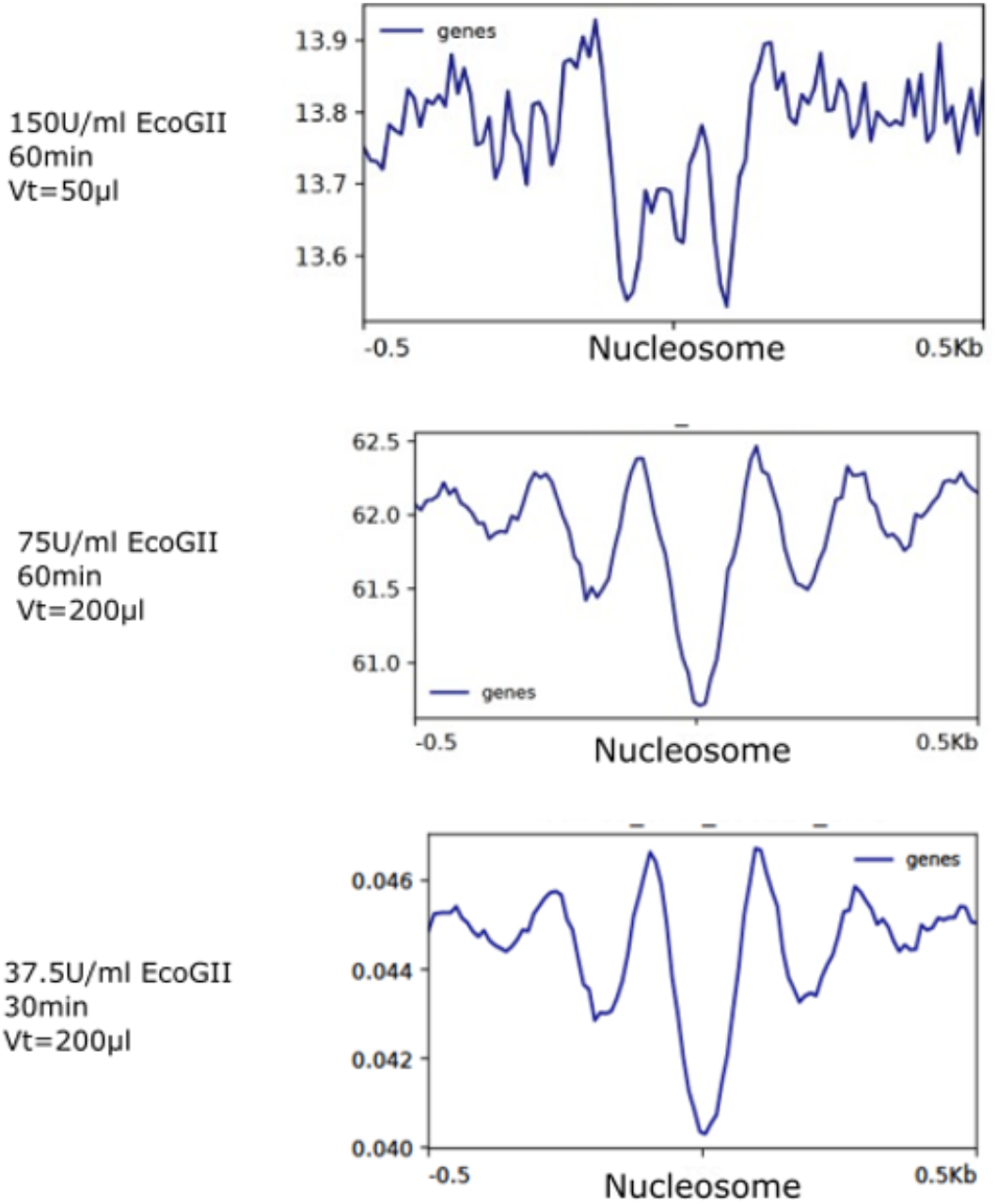
Decreasing EcoGII concentration and incubation time improves nucleosome resolution of SAM-seq. Vt indicates reaction volume.

**Supplementary Figure 7.**
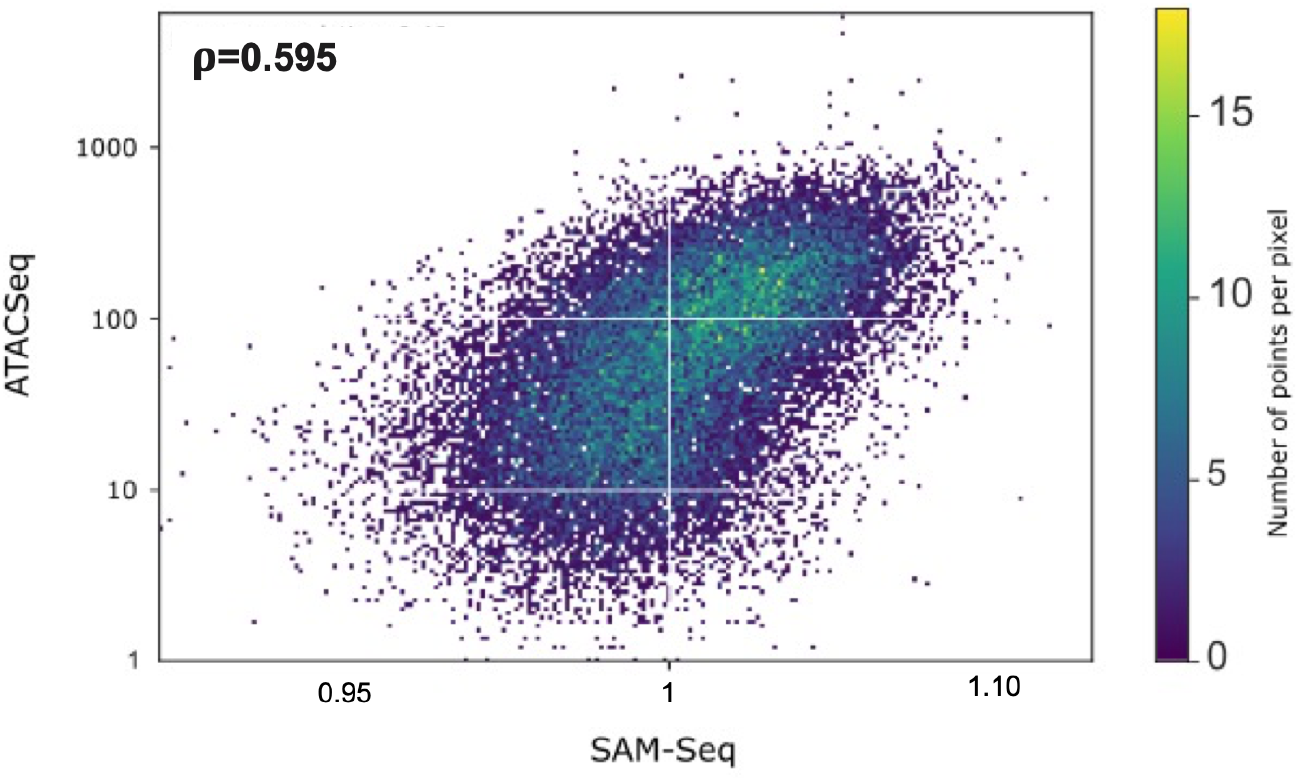
Pairwise comparison of accessibility between SAM-Seq and ATAC-Seq over 100bp regions upstream of TSS of Arabidopsis genes.

**Supplementary Figure 8.**
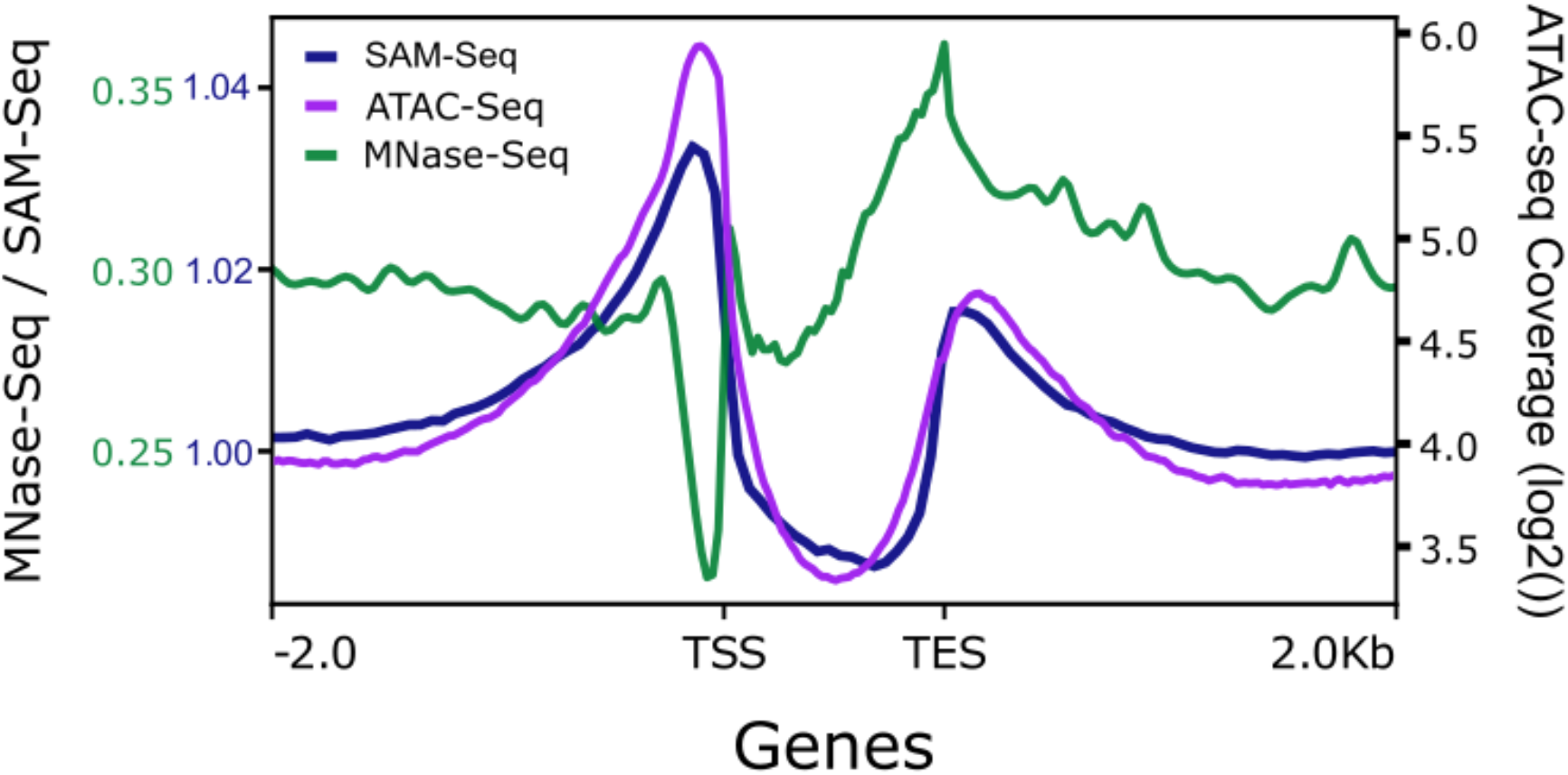
Metaplot of MNase-seq (Lyons et al. 2017), ATAC-Seq (Lu and al. 2017), and SAM-Seq accessibility over Arabidopsis genes.

**Supplementary Figure 9.**
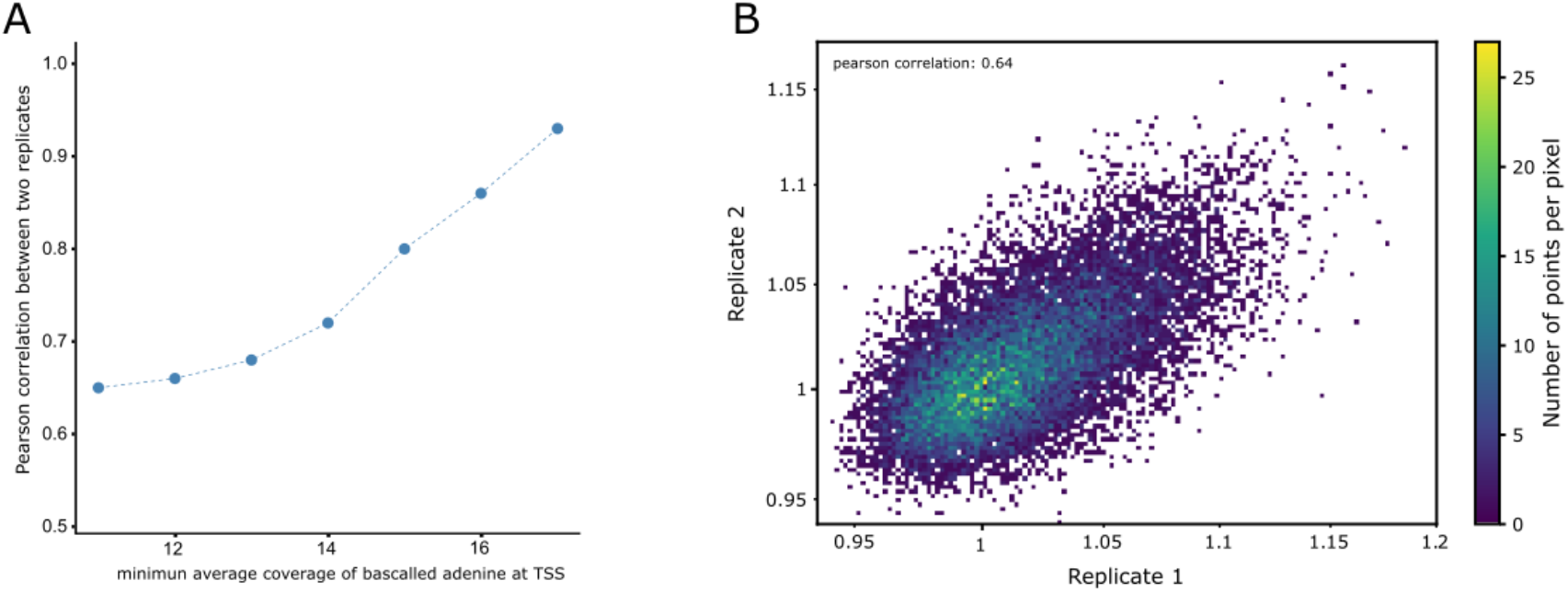
Pairwise comparison of accessibility at TSS of genes between two SAM-Seq Arabidopsis replicates. **A**. Pearson correlation between replicates of SAM-seq accessibility at TSS of genes is shown for different minimum average coverage of base-called adenine. **B**. Scatter plot showing normalized accessibility at TSS of genes for two SAM-seq replicates. TSSs with a minimum average coverage of 10 base-called adenines per strand were considered (first point in Panel A)

**Supplementary Figure 10.**
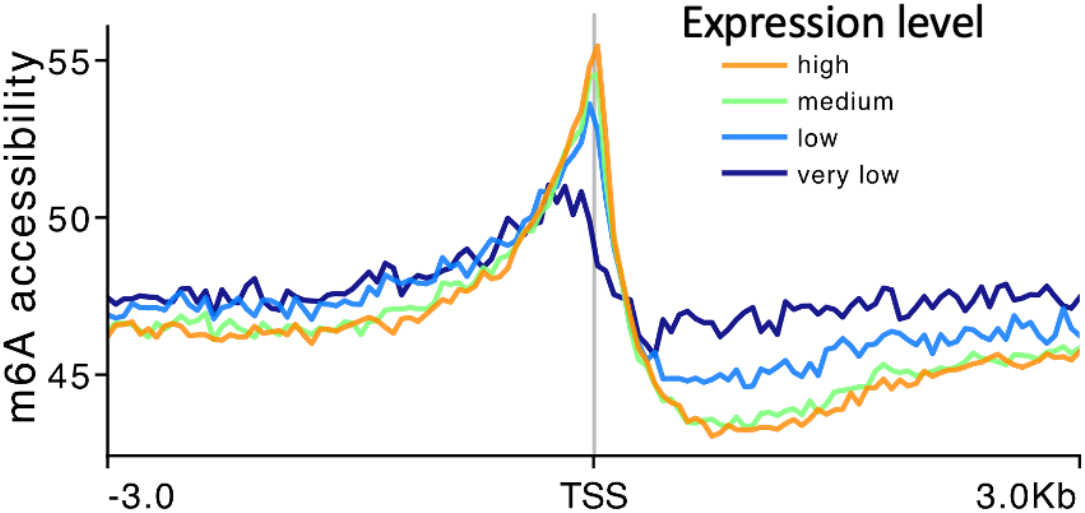
Metaplots of SAM-seq chromatin accessibility signal around transcriptional start sites (TSS) of Arabidopsis genes with different expression levels.

**Supplementary Figure 11.**
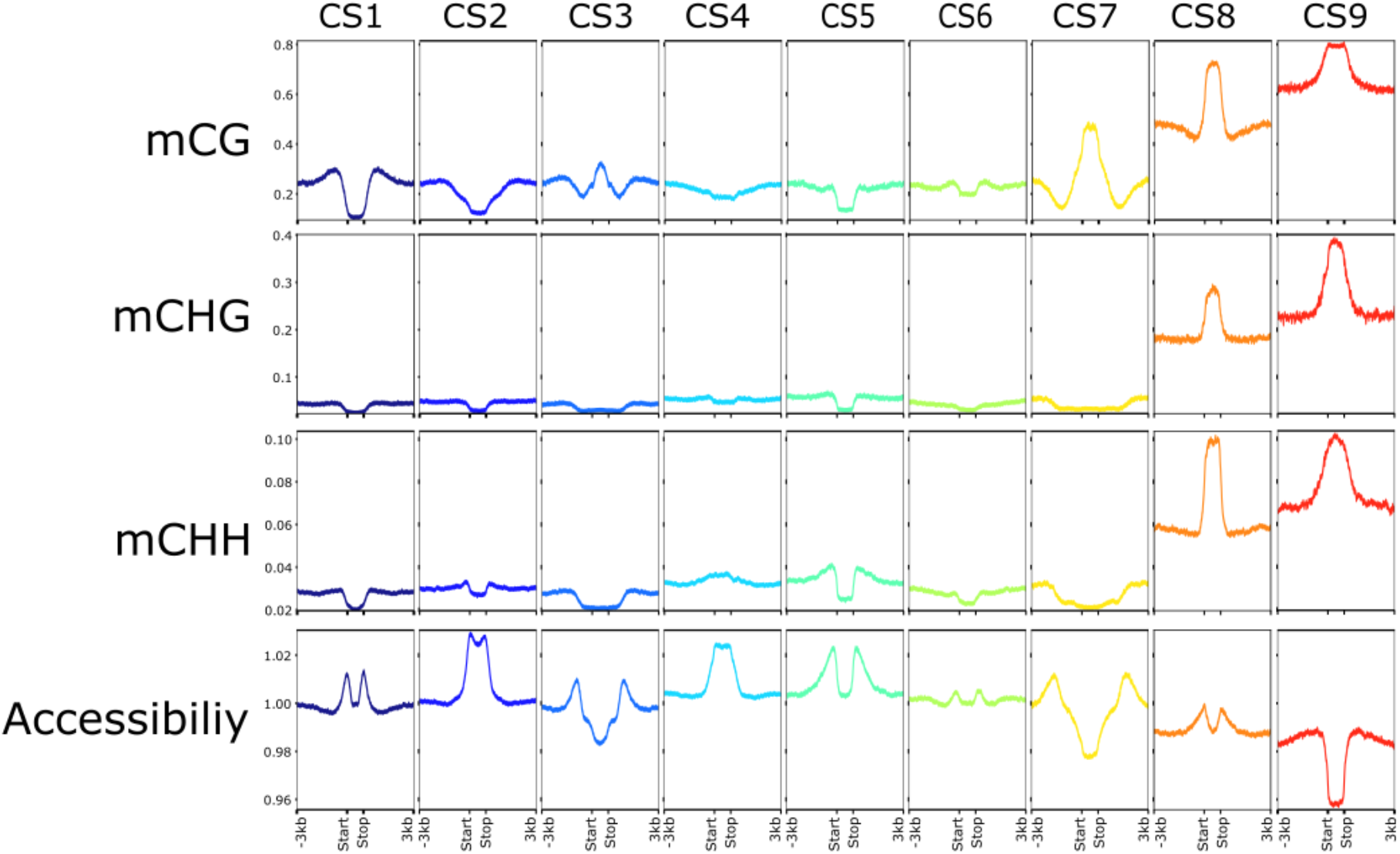
SAM-Seq accessibility and DNA methylation landscapes of Arabidopsis chromatin states. Accessibility, mCG, mCHG, and mCHH levels across nine chromatin states (CS) previously defined (Sequeira-Mendes *et al*., 2014). Each column corresponds to the indicated CS.

**Supplementary Figure 12.**
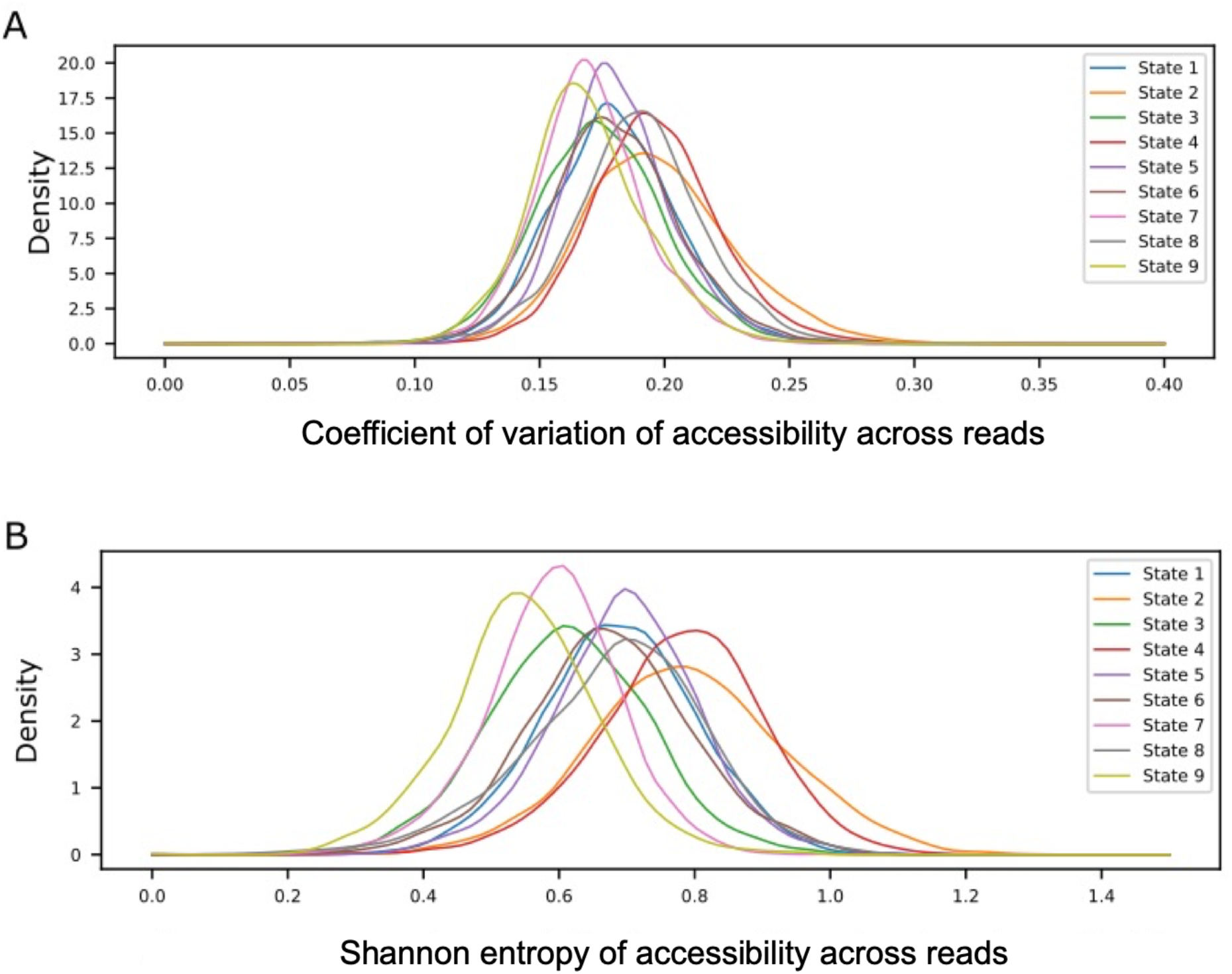
Heterogeneity of accessibility between fibres for the different Arabidopsis chromatin states**. A**. Coefficient of variation (CVs) of accessibility between fibres was calculated for each chromatin state feature. Density plots of distribution of CVs for the different chromatin states is shown. **B**. Shannon entropy of accessibility between fibres was calculated for each chromatin state feature. Density plots of distribution of Shannon entropy for the different chromatin states is shown.

**Supplementary Figure 13.**
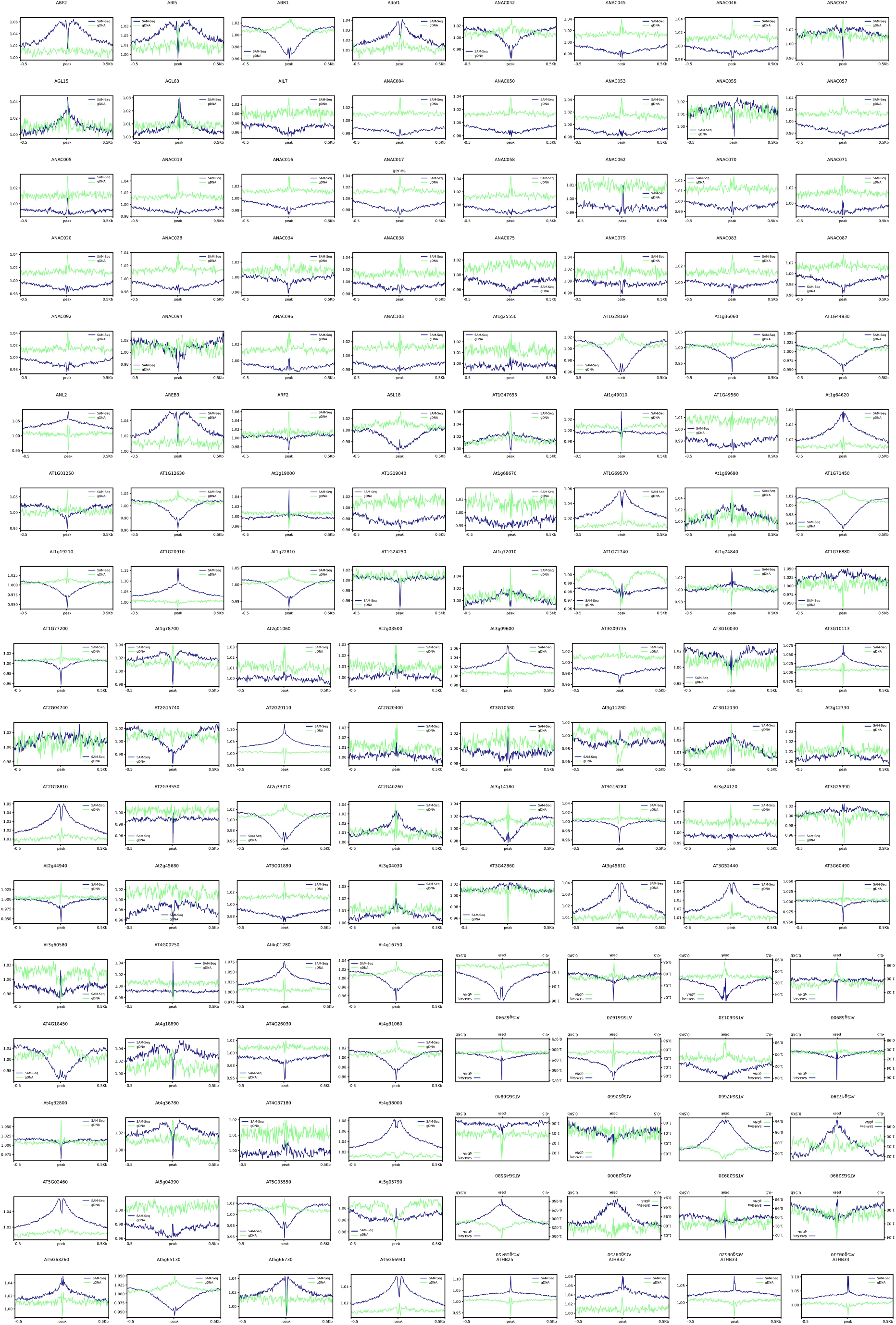
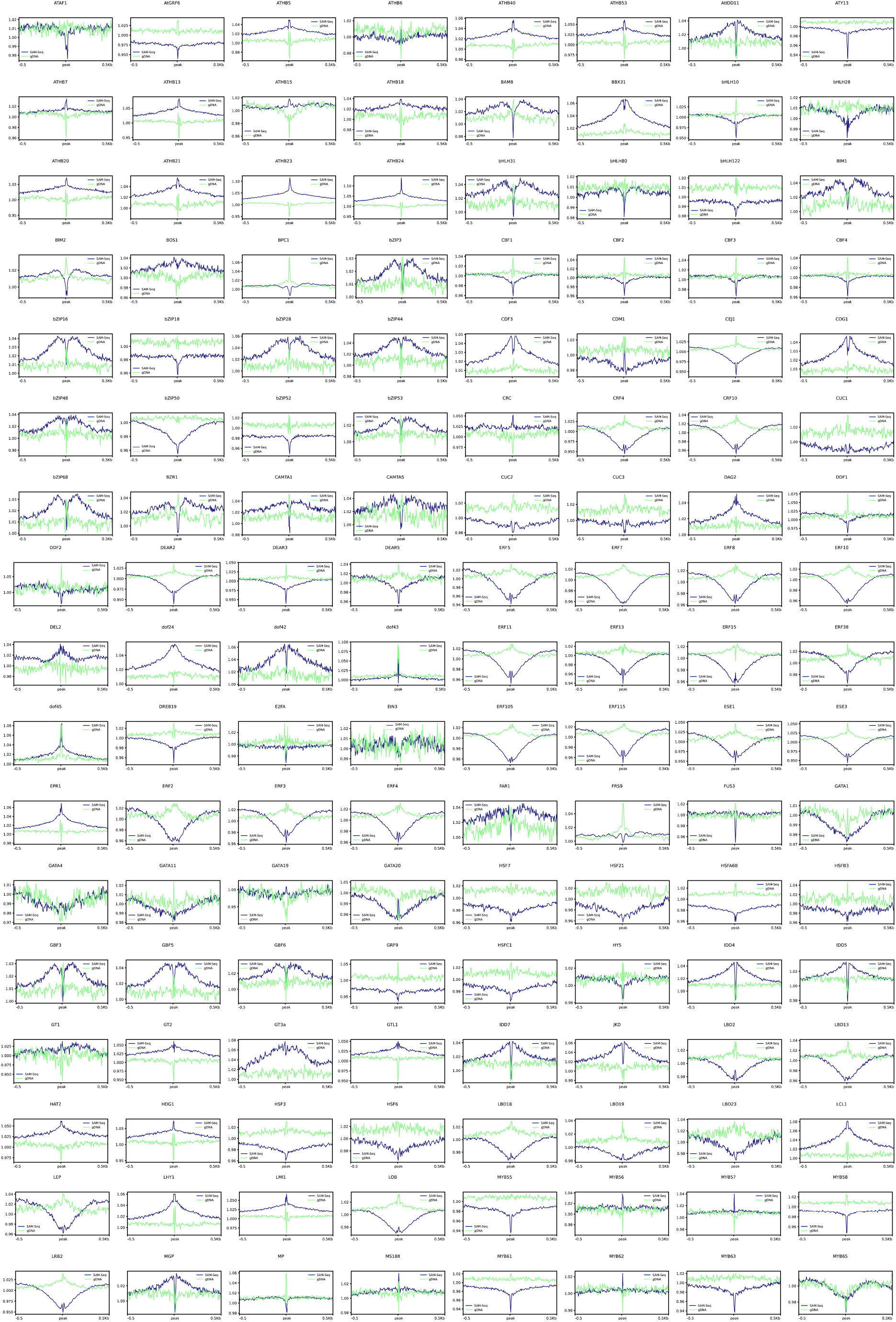
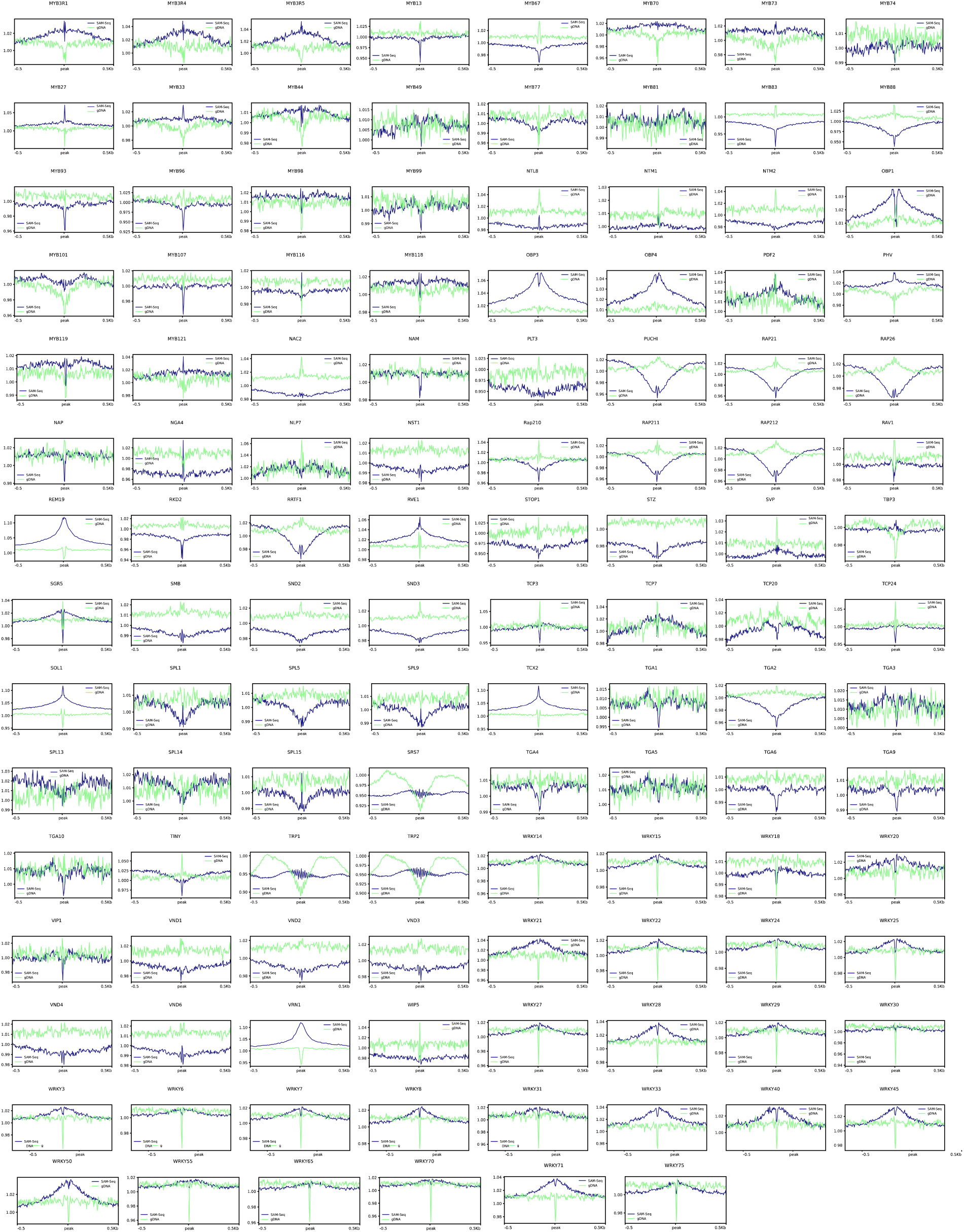
SAM-seq accessibility around 390 putative Arabidopsis TFBSs identified by DAP-seq. Metaplots of SAM-seq accessibility (SAM-seq), as well as 6mA/A levels from gDNA (gDNA) are shown around TFBSs previously detected (O’Malley *et al.,* 2016).

**Supplementary Figure 14.**
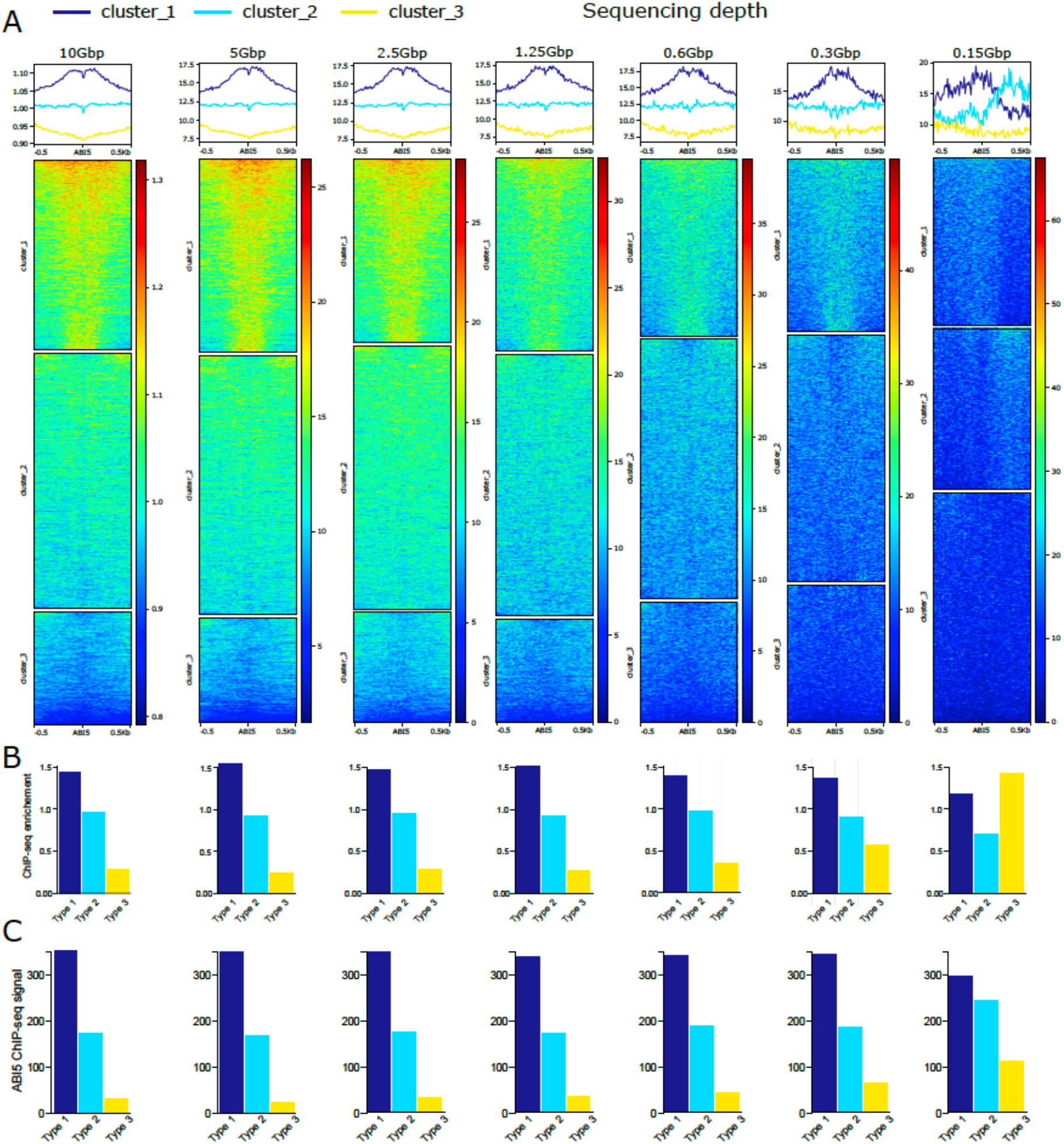
SAM-Seq footprinting pinpoints sites bound by Arabidopsis TFs in vivo at different sequencing depth**. A** SAM-seq experiment (10Gbp of sequencing depth) was sub-sampled progressively till reaching 0.15Gb. For each sub-sample, metaplot, heatmaps, and clustering of SAM-Seq chromatin accessibility around potential ABI5 binding site determined by DAP-seq were calculated. **B**. Enrichments of ABI5 Chip-Seq peaks at type 1, 2, or 3 potential ABI5 binding site. **C**. ABI5 Chip-Seq peaks signal over type 1, 2, or 3 potential ABI5 binding sites.

**Supplementary Figure 15.**
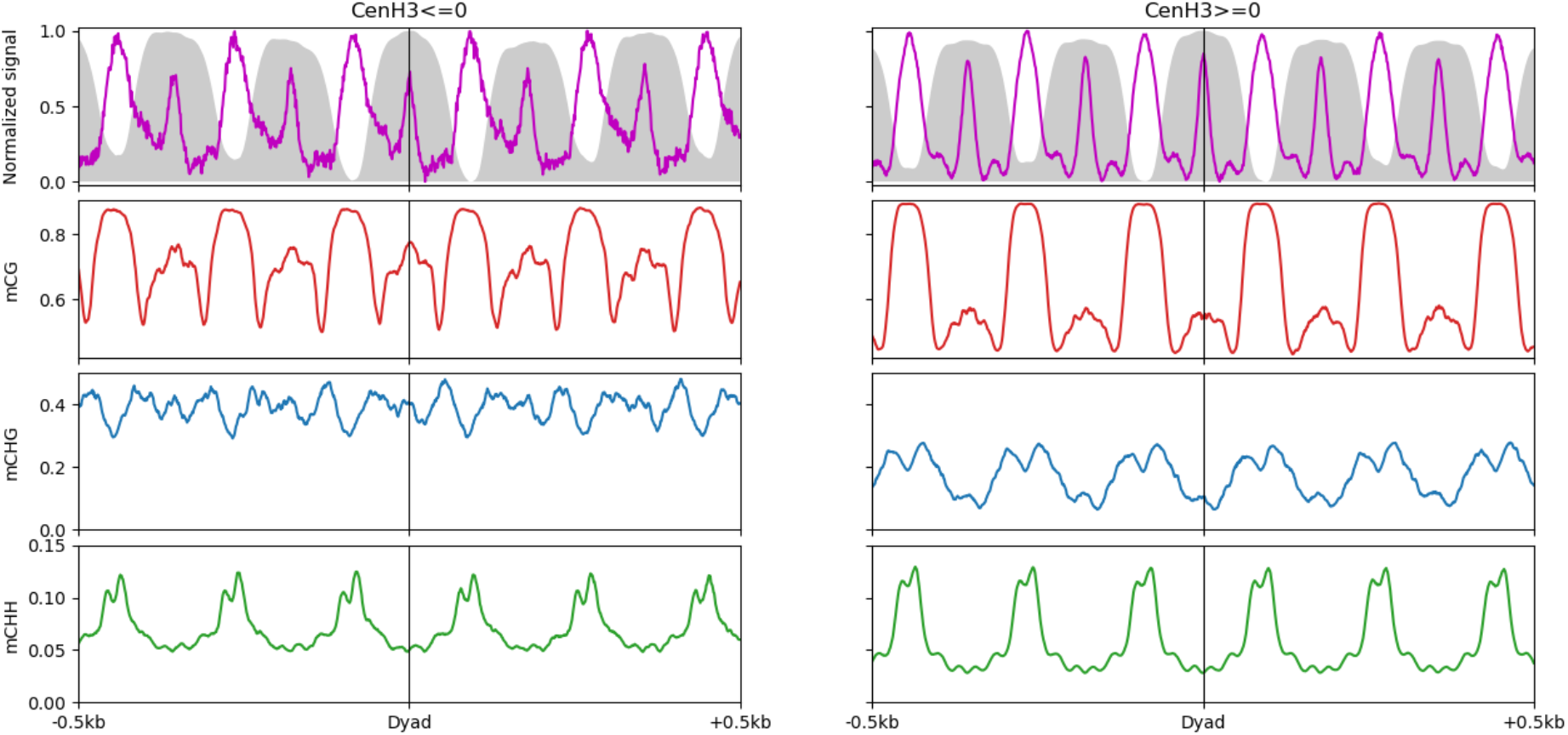
Accessibility and DNA methylation over nucleosomes spanning Arabidopsis centromeric repeats. Metaplots showing MNase (grey area), SAM-Seq accessibility (purple line), mCG (red), mCHG (blue) and mCHH (green) over well-positioned nucleosomes at centromeric repeats with high or low CENH3 levels.

**Supplementary Figure 16.**
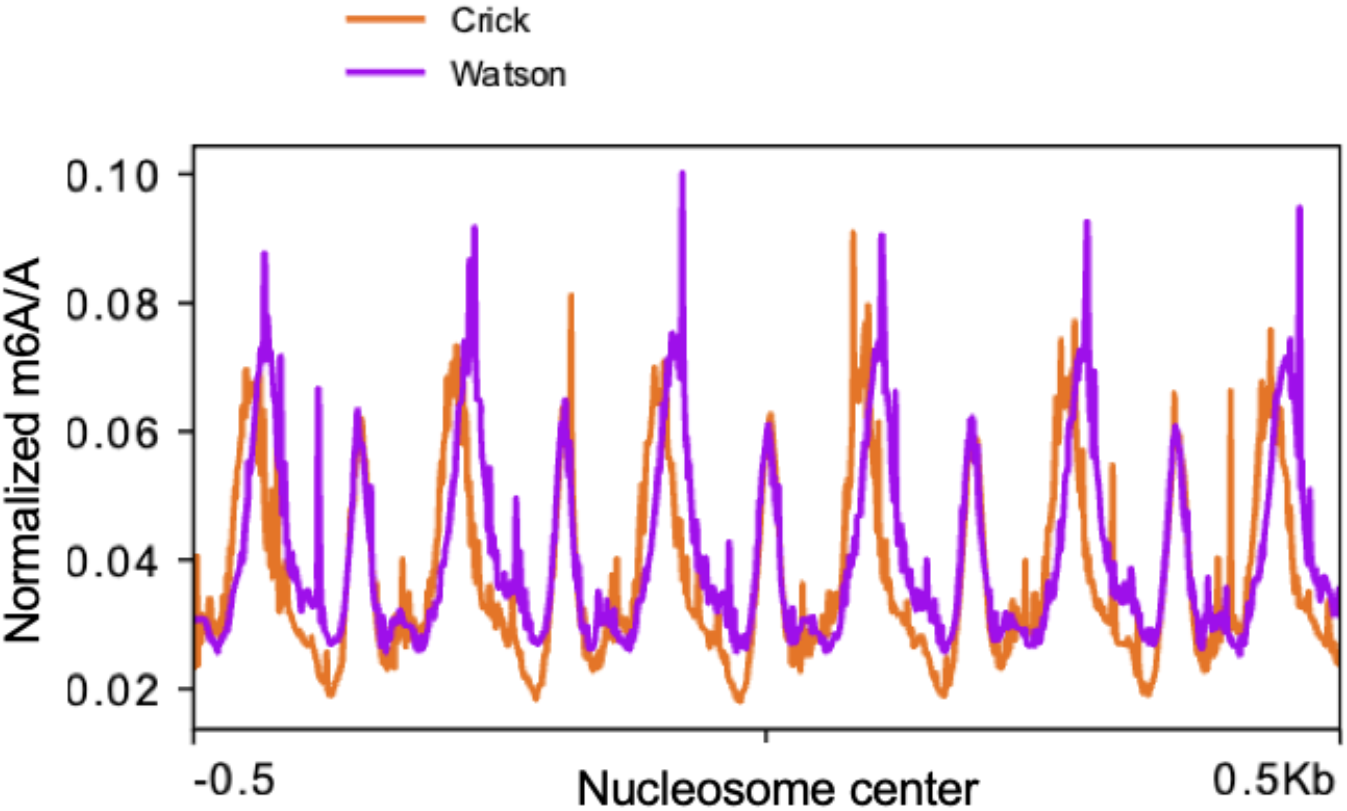
Metaplot of strand-specific SAM-seq accessibility around well-positioned Arabidopsis centromeric nucleosomes. Normalized SAM-Seq accessibility signal in the Watson (purple) and Crick (orange) strands are shown.

**Supplementary Figure 17.**
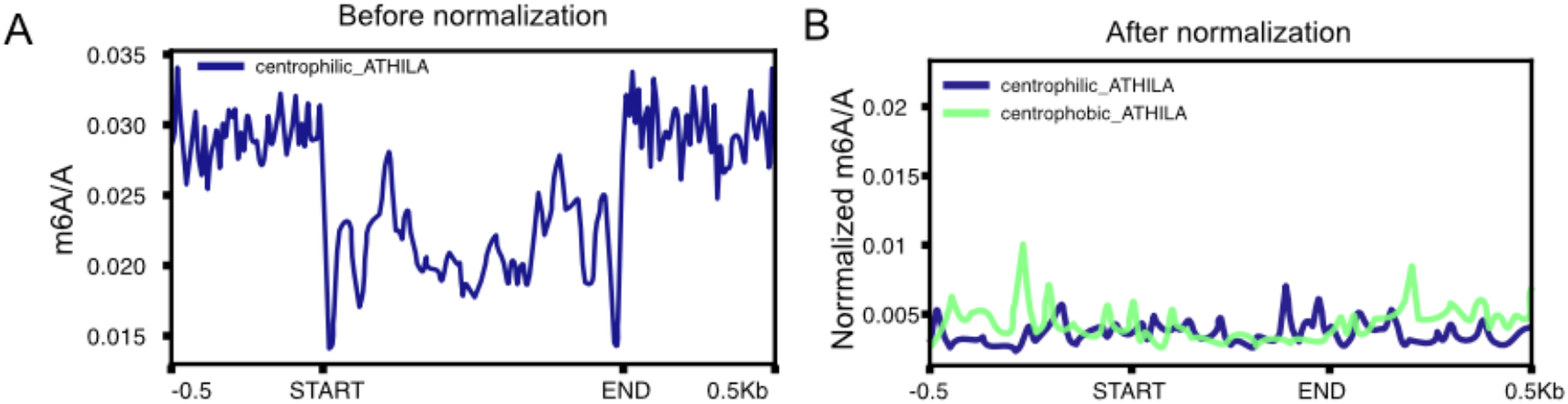
Metaplot of SAM-Seq accessibility signal over Arabidopsis centrophilic *ATHILA* LTR-retrotransposon. **A**. SAM-Seq accessibility signal at centrophilic *ATHILAs* before normalization. **B**. SAM-Seq accessibility signal over centrophilic and centrophopic *ATHILAs* after normalization.

**Supplementary Figure 18.**
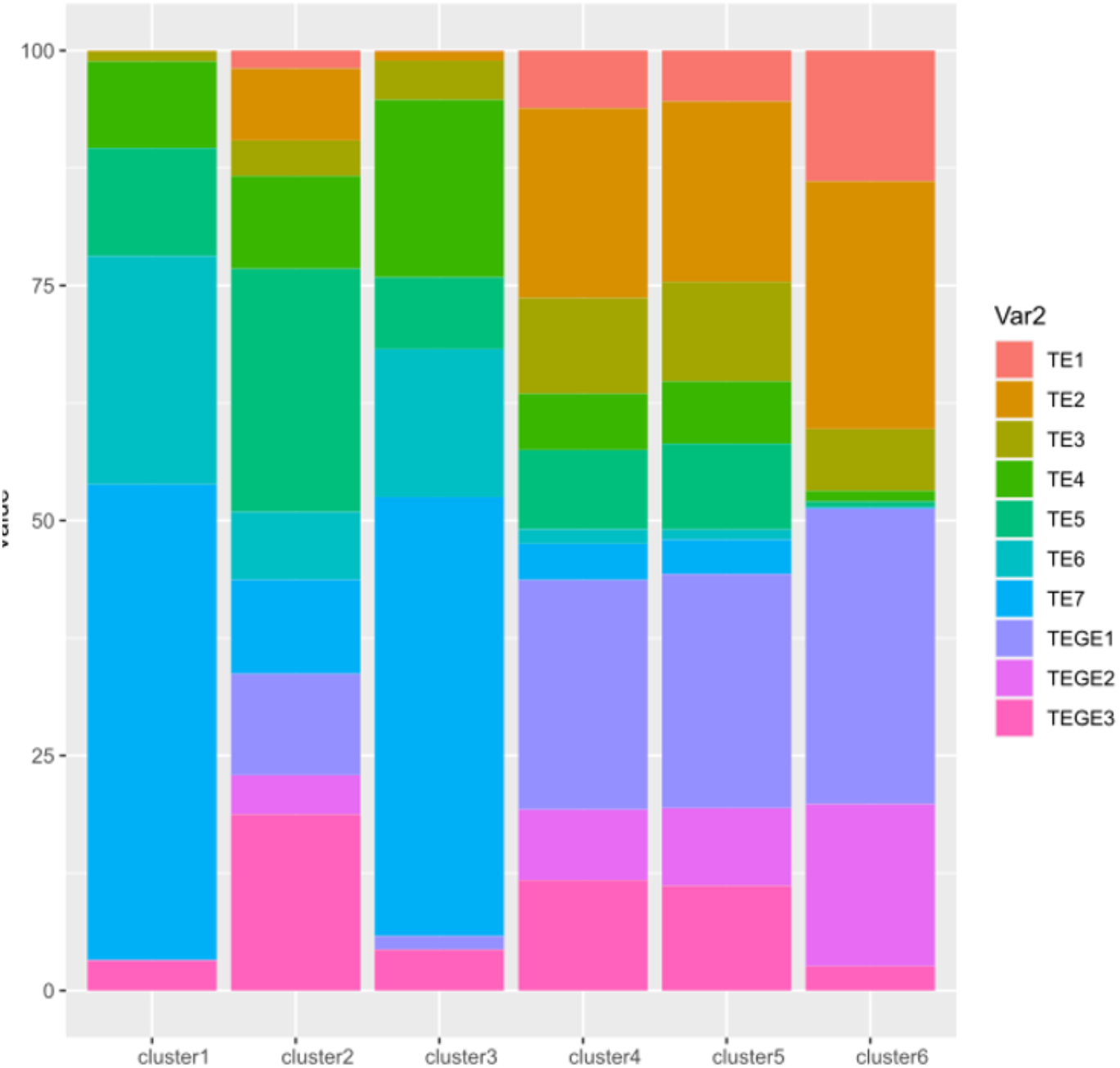
Overlap between Arabidopsis TE clusters identified by SAM-seq and those obtained based on multiple chromatin marks (Hisanaga *et al.,* 2023)

**Supplementary Figure 19.**
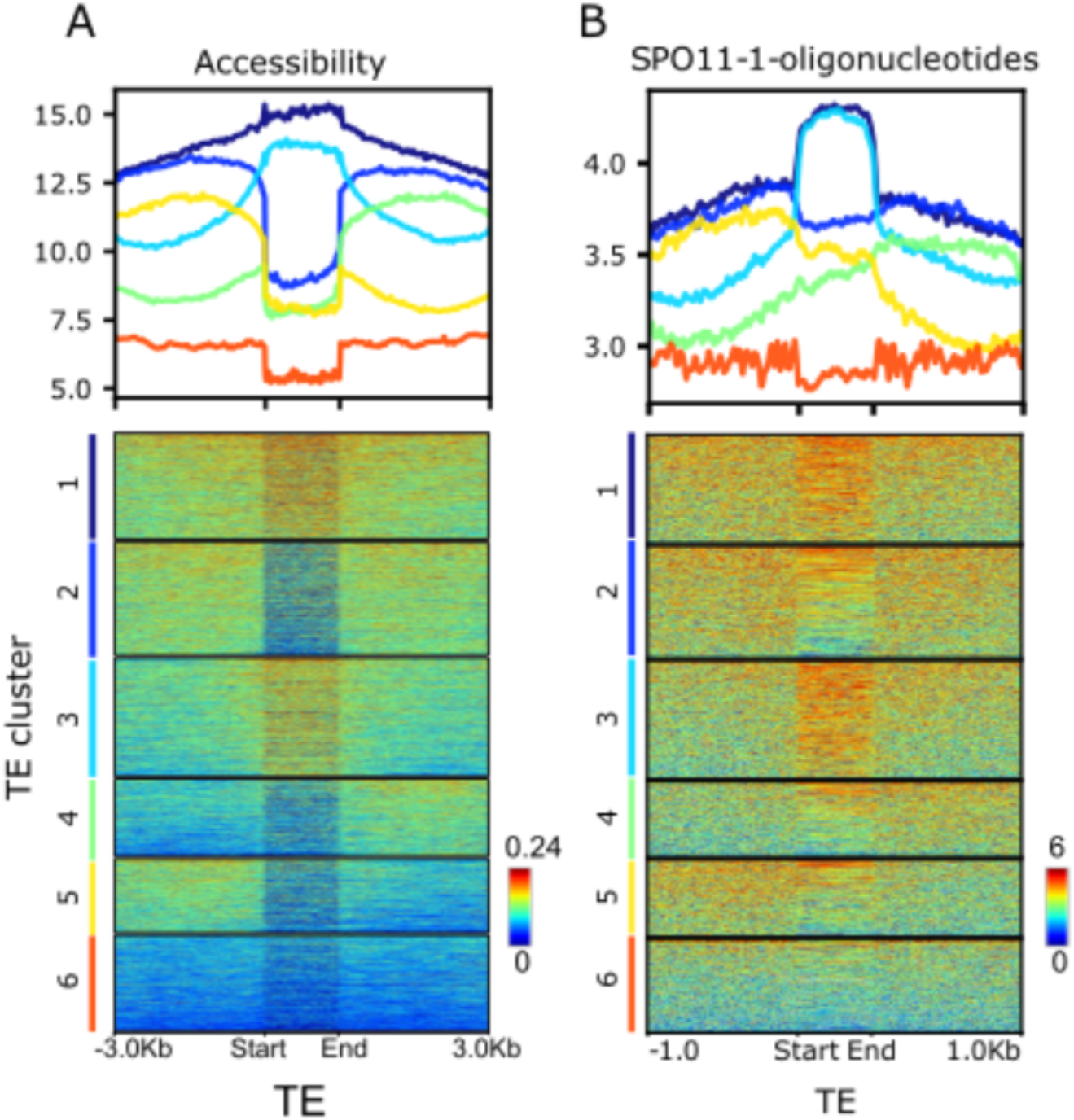
SPO11-1 oligos and SAM-seq accessibility profiling of Arabidopsis TEs**. A.** Metaplots and heatmaps for SAM-Seq accessibility across TEs in the Arabidopsis reference genomes. Clustering of TEs based on SAM-Seq accessibility identified six clusters. **B.** Metaplots and heatmaps for meiotic DSB initiation profiled by SPO11-1-oligonucleotide sequencing (Choi et al. 2018)

**Supplementary Figure 20.**
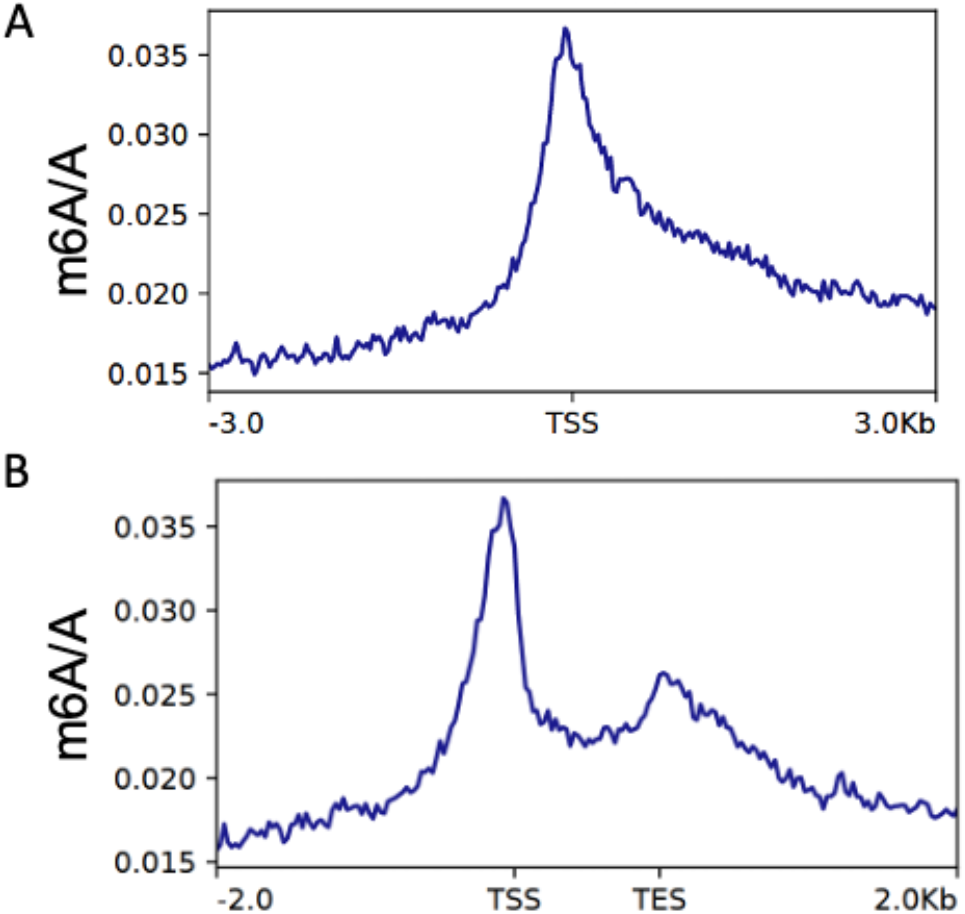
Metaplots of chromatin accessibility over genes for the non-reference Maize accession M52. **A**. Metaplots of SAM-seq chromatin accessibility signal around the Transcriptional Start Site (TSS) of genes. **B**. Metaplots of SAM-seq chromatin accessibility signal over genes. Transcriptional start and end sites (TSS and TES, respectively) are indicated.

**Supplementary Figure 21.**
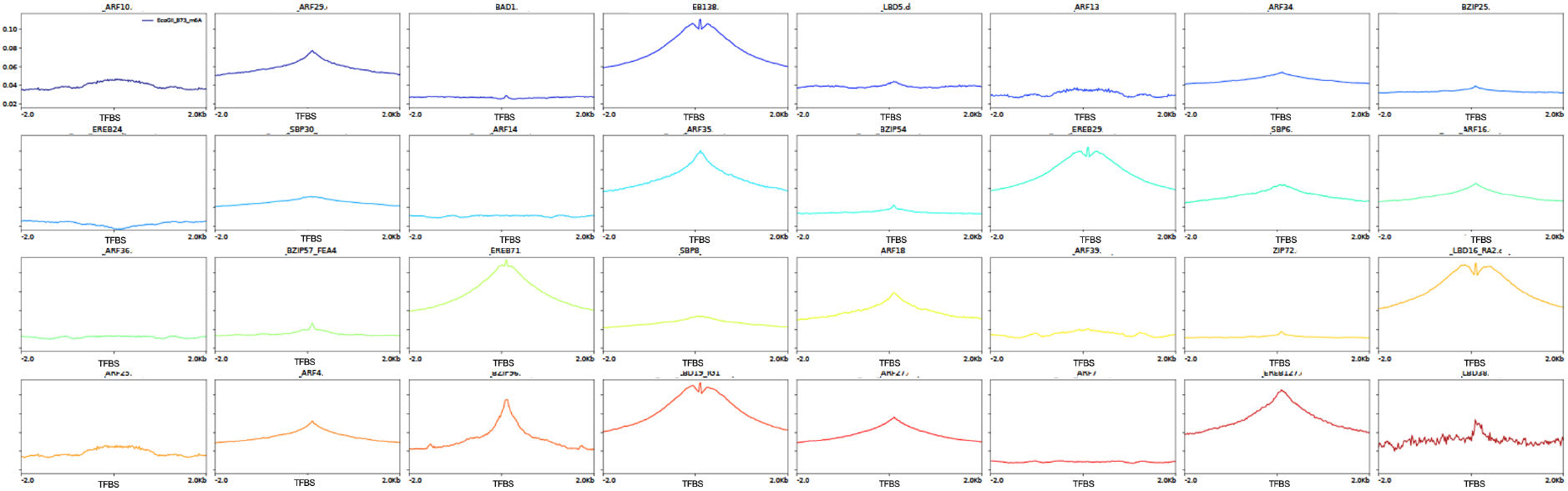
SAM-seq accessibility around 32 putative maize TFBSs identified by DAP-seq (O’Malley *et al.,* 2016 and Galli *et al*., 2018).

**Supplementary Table 1.**
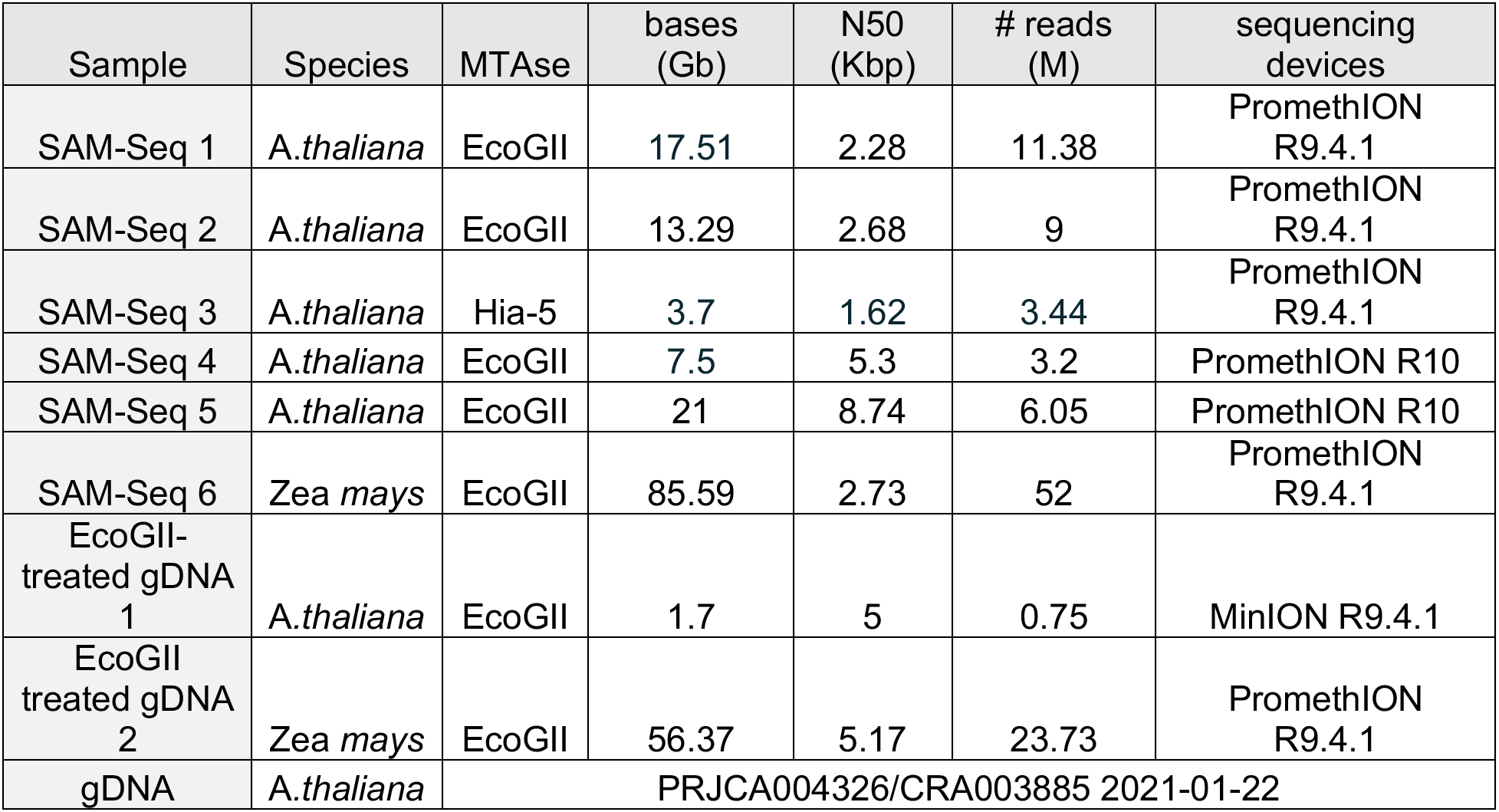
Sequencing summary metrics. Species, methyltransferase, sequencing depth, N50, number of reads and ONT chemistry are provided for each sequencing run.

| Sample | Species | MTase | bases (Gb) | N50 (Kbp) | # reads (M) | sequencing devices |
| --- | --- | --- | --- | --- | --- | --- |
| SAM-Seq 1 | *A.thaliana* | EcoGII | 17.51 | 2.28 | 11.38 | PromethION R9.4.1 |
| SAM-Seq 2 | *A.thaliana* | EcoGII | 13.29 | 2.68 | 9 | PromethION R9.4.1 |
| SAM-Seq 3 | *A.thaliana* | Hia-5 | 3.7 | 1.62 | 3.44 | PromethION R9.4.1 |
| SAM-Seq 4 | *A.thaliana* | EcoGII | 7.5 | 5.3 | 3.2 | PromethION R10 |
| SAM-Seq 5 | *A.thaliana* | EcoGII | 21 | 8.74 | 6.05 | PromethION R10 |
| SAM-Seq 6 | *Zea mays* | EcoGII | 85.59 | 2.73 | 52 | PromethION R9.4.1 |
| EcoGII-treated gDNA 1 | *A.thaliana* | EcoGII | 1.7 | 5 | 0.75 | MinION R9.4.1 |
| EcoGII-treated gDNA 2 | *Zea mays* | EcoGII | 56.37 | 5.17 | 23.73 | PromethION R9.4.1 |
| gDNA | *A.thaliana* | PRJCA004326/CRA003885 2021-01-22 |  |  |  |  |

